# Additive and dominance gene effects described by a logistic function in concentration-limited metabolic pathways at steady state out of equilibrium

**DOI:** 10.64898/2026.10.22.761079

**Authors:** Augusto Melgarejo, Alejandro R. Roig, B. Sofía Ramírez, Aníbal R. Lodeiro

## Abstract

Additive (*a*) and dominance (*d*) gene effects are key to phenotype manifestation. We investigated the relationship between these effects and gene-encoded enzyme rates in an *n*-step linear metabolic pathway with mass limitation at stationary state out of equilibrium, where internal flux depended on internal enzyme concentrations (*E*_*i*_; *i* = 1 to *n*) and external flux on input/output enzyme/transporter concentrations (*E*_*e*_). Using coupled reversible Michaelis-Menten (MM) equations for *E*_*i*_ rates and irreverisble MM for *E*_*e*_ rates we found that stationary-state metabolite concentration (*S*_*i*_) was a logistic function of *E*_*i*_. Its inflection point 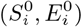 depended on the total mass of the system (Σ*S*_*i*_) and on *E*_*e*_ saturation, shifting from negative to positive values as reaction kinetics transitioned from first to mixed or zero order. Moreover, *Σ S*_*i*_ and *E*_*e*_ saturation explained negative, partial, and complete dominance in this metabolic system, provided phenotypic value of trait *T* is linearly dependent on *S*_*i*_ at certain sink. The system was unstable when approaching closure (*E*_*e*_ *→* 0). In this condition, asymptotic 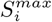 equaled equilibrium concentration, being equilibrium displacement the driving force of flux. In pathways with *n* > 1 reaction steps there was a change of regime in early reactions when increments in *E*_*n*_ passed 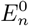. This change of regime led to peaks raising at some *S*_*i*_ previous to *S*_*n*_ and *S*_*n*+1_ that were modeled by adding a Gaussian component to the logistic equation. Therefore, if *T* is determined by some of these *S*_*i*_, overdominance and dominance reversals may be explained. We obtained a good fit when applied this equation to *E*_*i*_ and *S*_*i*_ data obtained from silicon-cell models developed with specific kinetic equations and experimentally determined kinetic parameters. In addition, we applied this framework to experimental data on amylose accumulation in starch granules catalyzed by granule-bound starch synthase I, and found that the logistic function fitted data from six plant species. This framework may contribute a partial explanation of the non-linearity between genotype and phenotype from the metabolic standpoint, which may be useful in the future for plant or animal breeding technologies.

## I. INTRODUCTION

The connection between genotype and phenotype, known as genetic architecture, is a cornerstone of genetic analysis. Classical models elegantly dissect this relationship into core components—additive gene effects, dominance intragenic effects, epistatic intergenic effects, environmental effects, as well as the interactions between all them. However, the interpretation of dominance depends on whether a trait is linked to fitness, such as a life-history trait, or if it is a non-fitness-related or morphological trait [1]. For life-history traits, dominance measures how closely a hybrid’s viability aligns with that of each homozygous parent, while for morphological traits, it reflects the degree to which a hybrid’s phenotype resembles one parental extreme over the other [2]. For applications to plant or animal breeding, where the evolutionary perspective is less important, dominance is considered in the Mendelian, morphological sense. Here we will adopt this point of view, where the additive gene effect (*a*) represents the half-difference in phenotypic value between two contrasting homozygous parents *P*_1_ and *P*_2_ and dominance (*d*), the deviation of the hybrid progeny *H* from the midpoint *m* between the parents without regard to the fitness of these genotypes. Thus, *a* and *d* may be written as [3]

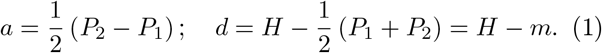

In agricultural research, numerous important traits are expressed in quantitative units and their inheritance may be investigated in terms of *a* and *d*. Consequently, these effects become subjects of statistical analysis in understanding quantitative traits and to find the *loci* (QTL) responsible for these traits in the DNA [3]. However, the connection between the statistical significance of parameters *a* and *d* and their underlying molecular mechanisms remains poorly understood. This is particularly challenging in the case of *d*, as it resists straightforward mechanistic interpretation due to the complex expression and integration of gene products within metabolic networks [4–6].

The simplest molecular approach to understand *d* at the level of the metabolic pathway is by assuming constitutive genes encoding monomeric enzymes that catalyze a linear set of monosubstrate chemical reactions, whose final product is responsible for phenotype. For instance, let us suppose a given gene *G*, for which *a* and *d* effects may be though to depend on how gene dose determines the encoded enzyme concentration *E*, and how much *E* is required for the full expression of a trait *T*. In this simple frame, we may also suppose that *G* possesses two alleles, in such a way that allele *G* encodes the fully active enzyme while allele *g* encodes a defective form. The simplest case of dominance in a *Gg* hybrid derived from *GG* × *gg* cross is no dominance (*d* = 0) in which the hybrid *Gg*, having 50% *E*, shows 50% *T* of the homozygous parent *GG* (eq. 1). However, since gene-encoded enzymes are embedded in metabolic networks, its placement and interactions with other enzymes condition gene effects, even for this simple idealization. Furthermore, cells—where metabolic pathways occur—function as open systems at steady-state far from equilibrium. They maintain a relatively constant total concentration of metabolites, constrained by factors such as cell size, reaction rates, and osmotic homeostasis. Therefore, the relationship between *E* and *T* is non-linear and, depending on the metabolic structure and dynamics, a range of situations in which *d* ≠ 0 occur, being complete dominance (*d* = *a*) the known case of Mendel’s traits [7, 8]. Many attempts to mechanistically explain these different relationships between allele dose, enzyme concentration, trait expression, and dominance were addressed in terms of metabolic control theory [9–11]. In these studies, emphasis was put in explaining how *E* determines the metabolic flux *J* through the pathway of interest. However, most of gene effects determining *T* should be better explained in terms of stationary-state metabolite concentrations *S* rather than in terms of *J* [12, 13], although it is evident that both magnitudes are closely related (here “metabolite” is used in a broad sense, including signaling molecules, polymers, etc.). In addition, these methods seldom took into account the limitation to metabolite concentrations imposed by cell size, and in many instances they worked at substantially low metabolite concentrations in order to simplify the mathematical treatment by approaching first-order kinetics. Therefore, a description of metabolic output in response to changes in the concentration of an enzyme catalyzing a specific pathway step, capable of accounting for the constraints imposed by the total metabolite concentration within the cell and the cell’s inherent nature as an open system in a steady-state far from equilibrium, may contribute to a better understanding of additive and dominance gene effects.

In addition, a new demand for such methods is raising. Increasing information about biochemical systems is being provided at unprecedent rate by high throughput technologies such as next-generation DNA and RNA sequencing, mass spectrometry-based proteomics and metabolomics, and machine learning data analysis [14– 16]. Nevertheless, enzyme and metabolite datasets were seldom correlated with each other in a single study [17]. An opportunity to correlate allelic variants with enzyme and metabolite concentrations in large datasets is now being provided by microfluidics-based approaches such as high-throughput microfluidic enzyme kinetics and deep mutational scanning [18–20]. However, to our knowledge, these methods were used with isolated enzymes or cell-free systems, and their purpose was mostly to enhance enzymes for biotechnological applications, but it seems that it is only a matter of time for they become applied to broader fields such as plant and animal breeding, where *a* and *d* play key roles.

In this work, we modeled a linear metabolic pathway occurring in a cell (or cell compartment) of finite size at steady-state far from equilibrium. These constraints revealed that the *S*_*i*_(*E*_*i*_) relationship for the *i*-th step may be analyzed with a logistic function, regardless of the number of steps. By assuming that the genotypes *G*_*i*_*G*_*i*_, *G*_*i*_*g*_*i*_, and *g*_*i*_*g*_*i*_ for the gene encoding an enzyme produce *E*_*i*_ concentrations distributed along the x-axis of the logistic curve, we obtained a novel description of *a* and *d* gene effects, which we further validated using *in silico* models and experimental data.

## II. A GENERAL PERSPECTIVE

Stability of open systems far from equilibrium, which exchange matter and energy with the surroundings in continuous flow, is achieved when fluxes get constancy at stationary state for which the equilibrium state is a particular case occurring in the closed system. Based on that, we can identify a state out of equilibrium through a vector of the form

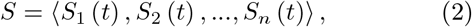

where *S*_*i*_ (*t*) represents the concentration of the *i*th metabolite of the pathway of interest at time *t*.

For simplicity, we will focus on a linear metabolic pathway with *n* monosubstrate reactions at steady concentrations *S*_*i*_ *⇌ S*_*i*+1_ catalyzed by monomeric enzymes at concentrations *E*_*i*_ (where *i* = 1…*n*) [21]

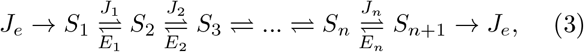

where *J*_*e*_ is the external flux. This pathway can be conceptualized either as a truncated segment of the global metabolic network or as the terminal enzymatic steps preceding product efflux. Under either interpretation, the flux *J*_*e*_ denotes the enzyme-catalyzed final step or the transporter-dependent export of mass toward an external sink. We contend that a kinetic description of *J*_*e*_ based on irreversible Michaelis–Menten formalism is physiologically more realistic than either first-order or zero-order alternatives. Unlike first-order kinetics, which assumes a linear dependence on substrate concentration and fails to capture saturation effects, or zero-order kinetics, which neglects the regulatory influence of substrate availability, the Michaelis–Menten equation inherently accounts for the finite catalytic capacity of the enzyme/transporter and the progressive occupancy of its active site as substrate concentration increases. This provides a more faithful representation of the underlying enzymatic mechanism, particularly under conditions approaching enzyme/transporter saturation. The canonical form of this rate law is:

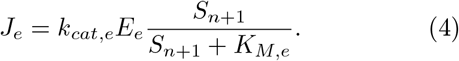

where *k*_*cat,e*_ denotes the catalytic constant (turnover number), *E*_*e*_ is the concentration of the exporting enzyme or transporter, *S*_*n*+1_ is the concentration of the terminal product, and *K*_*M,e*_ represents the corresponding Michaelis constant. In addition, to get the stationary state condition we set the input flux as equal to the output flux, so that both entry and exit reactions are considered identical. In turn, kinetics of each internal flow *J*_*i*_ was represented by the reversible Michaelis-Menten relationship

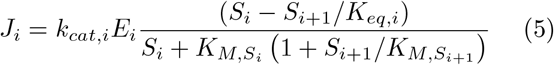

where *k*_*cat,i*_, *E*_*i*_, *S*_*i*_ (or *S*_*i*+1_), and 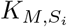 (or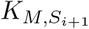) represent the same magnitudes as in the external flux for the internal *i*-th step (*i* = 1 to *n*) and *K*_*eq,i*_ is the equilibrium constant of the corresponding reaction. The rationale of choosing reversible kinetics for the internal flux was for considering reaction product effects in the internal reactions. In the following, we used as kinetic parameters: 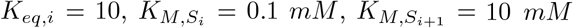, *K*_*M,e*_ = 1 *mM* , *k*_*cat,i*_ = 10000 *min*^*−*1^, and *k*_*cat,e*_ = 10000 *min*^*−*1^.

In light of the above considerations, we neither require further assumptions on the relative concentrations of metabolites at the input and output, nor to adjust metabolite concentrations to a situation far from enzyme saturation [11]. Instead, in this model the relationships between *E*_*i*_ and *S*_*i*_ are alike those reported in living cells [22, 23], and the external flux kinetics may span either first order (*S*_2_ *≪ K*_*M,e*_), mixed order (*S*_2_ *≫ K*_*M,e*_), or zero order (*S*_2_ *≃ K*_*M,e*_) [24].

The dynamics of the state represented by relation (2) may be studied through a system of coupled differential equations (DE), which in generic form is expressed as

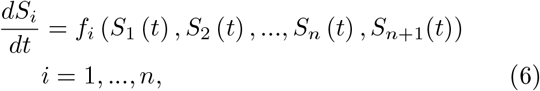

where the functions *f*_*i*_ (*S*_1_ (*t*) , *S*_2_ (*t*) , …, *S*_*n*_ (*t*) , *S*_*n*+1_(*t*)) are written in terms of the internal and external flows (4) and (5).

System (6) considers the stoichiometry of the metabolic reactions that take place during the process. In particular, here we will be interested in the stationary states, for which *f*_*i*_(*S*_1_, …, *S*_*n*+1_) = 0.

## III. ANALYSIS OF A PATHWAY WITH A SINGLE STEP

We will start with the analysis of a pathway composed by a single reaction step in an effort to identify and understand the parameters that affect the relationship between *E*_*i*_ and *S*_*i*_ in an open system at stationary state far from equilibrium. Let us assume that metabolite 1 enters the pathway at *S*_1_ concentration, where it is transformed into metabolite 2 at *S*_2_ concentration in a reaction catalyzed by enzyme 1 at *E*_1_ concentration, being the magnitude of trait *T* linearly dependent on *S*_2_ in a given sink

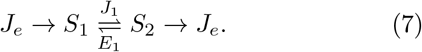

Since this reaction is unimolecular, *J*_*e*_ and *J*_1_ relate to *S*_1_ and *S*_2_ as follows:

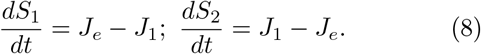

At the steady-state out of equilibrium, necessarily *J*_*e*_ = *J*_1_ and, provided no gain or loss of matter occurs, the total mass of the system must remain constant. Therefore, to analytically determine the steady-state *S*_1_ and *S*_2_, we begun by equating (4) and (5) taking into account the conservation of mass (Σ*S*_*i*_ = *constant*). With this procedure, we analyzed the change in *S*_*i*_ in response to changes in *E*_1_ as the relationship between *dS*_*i*_*/dE*_1_ and *S*_*i*_ (for details, see Supplementary Material)

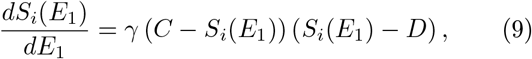

with *C, D ∈* ℝ. Its solution corresponds to the logistic function, given by

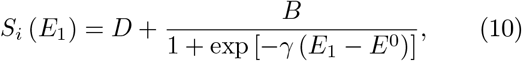

where *B* = *D/*(*C* − 1), with *C* ≠ 1. Before proceeding further, we find it instructive to express the logistic function (10) in an alternative form

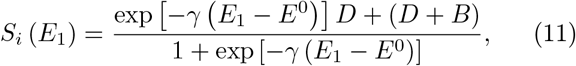

which may be useful for assigning a clearer biological or kinetic meaning to each of the constitutive constants appearing in the model. By replacing 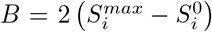 and 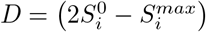 we obtain:

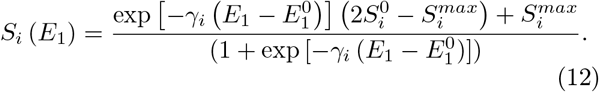

In this reformulation, 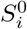 and 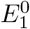are *S*_*i*_ and *E*_1_ at the inflection point respectively for a given value of *E*_*e*_, *γ*_*i*_ is a measure of the rate of change of *S*_*i*_ per unit change in *E*_*i*_ at the inflection point, and 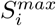 is the asymptotic *S*_*i*_ attained at highest *E*_1_. Hence, 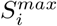 may be interpreted as the carrying capacity of the system [25], and *γ*_*i*_ as a measure of the rate at which the carrying capacity is reached.

Our analytical approach suggested that the logistic equation may represent the change in *S*_*i*_ in response to changes in *E*_1_. Hence, we inquired as to whether this equation might fit numerical solutions of the system represented by (4) and (5) at steady-state. To this end, we performed the calculations starting from a set of initial conditions *S*_*i*_ (0) where each metabolite had the same concentration at *t* = 0 of the calculation. Then, we run this model following (6) until a time interval Δ*t* over which all *S*_*i*_ remain constant, which we take as the stationary concentrations. Therefore, hereafter we will omit the time dependence (*t*) when referring to these stationary *S*_*i*_.

In a first approach we choose the set of *S*_*i*_ (0) = 0.1, 1.0, 3.0 and 10.0 *mM* and calculated stationary *S*_1_ or *S*_2_ as function of *E*_1_ [denoted *S*_*i*_ (*E*_1_)]. In addition, mass conservation led us to consider *nS*_*i*_(0) = Σ*S*_*i*_. With this procedure, we observed that, indeed, (*S*_*i*_, *E*_*i*_) followed continuous sigmoidal curves (Fig. 1 and Supplementary Fig. S3). As we will see later, equation (12) holds in general for the last step of all linear pathways tested here, regardless of the number of intermediates.

**Figure 1:**
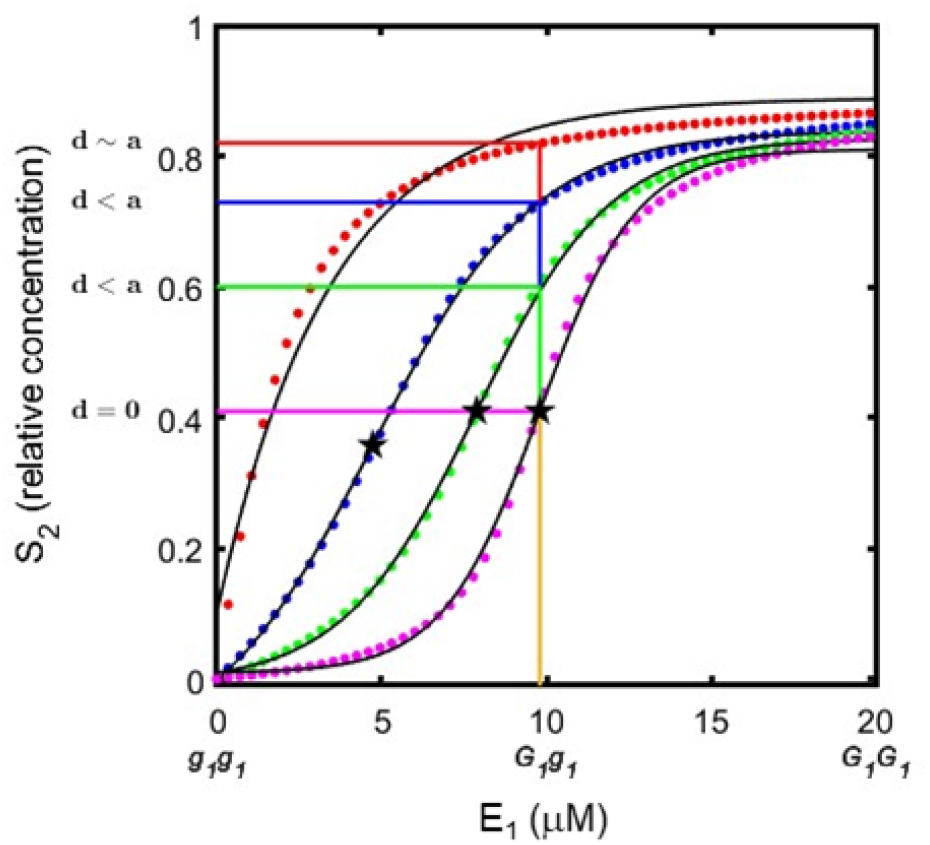
Product concentration (*S*_2_) in response to increases in active monomeric internal enzyme concentration (*E*_1_) in an open system at steady-state out of equilibrium (*E*_*e*_ = 10 *µM*). Total concentrations Σ*S*_*i*_ were: 0.2 *mM* (red), 2.0 *mM* (blue), 6.0 *mM* (green), and 20.0 *mM* (pink). Black lines are the fitting of data with eq. 12 (0.98 *≤ R*^2^ *≤* 0.99), the stars indicating the inflection point (for Σ*S*_*i*_ = 0.2 *mM* inflection point is not shown because in this condition it lies at the negative quadrant). Each *G*_1_ allele encodes one copy of fully active enzyme while *g*_1_ allele encodes a defective enzyme *−*which does not contribute to *E*_1_. Dominance for each Σ*S*_*i*_ is indicated at the *ordinate*. Relative *S*_2_ values are *S*_2_*/*(*S*_1_ + *S*_2_) to ease comparison among the different Σ*S*_*i*_. The corresponding curves for substrate (*S*_1_) are in Supplementary Fig. S3.

Assuming that enzyme 1 is encoded by gene 1 with two alleles, *G*_1_ and *g*_1_, such that the homozygous *G*_1_*G*_1_ produces twice as much *E*_1_ as *g*_1_*g*_1_, and the hybrid *G*_1_*g*_1_ has an *E*_1_ level midway between both homozygous, it follows that the degree of dominance can shift from no dominance to complete dominance depending on Σ*S*_*i*_ (Fig.1), which may be tissue-specific or vary along time at longer periods than considered for the analysis of steadystate. Moreover, owing to stochastic fluctuations in gene expression, protein folding, and degradation rates, the effective *E*_1_ associated with a given genotype can vary among individual cells [26, 27], all of which may contribute to environmental variance within a population. However, beyond this intra-specific variation, a more fundamental principle emerges when considering different taxons. While the relationship between *S*_2_ and *E*_1_ shown in Fig.1 follows a universal sigmoidal form for a given metabolic context, the absolute levels of enzyme activity achieved by the homozygous genotypes *G*_1_*G*_1_ and *g*_1_*g*_1_ are not fixed across organisms. Taxon-specific differences in regulatory baselines, genomic background, gene interactions, or small variations in protein sequences can reposition orthologous *G*_1_*G*_1_ and *g*_1_*g*_1_ homozygous along the *S*_2_(*E*_1_) curve, yielding dominance relationships that range from none to complete as a consequence of metabolic and evolutionary context [28–33].

In addition to Σ*S*_*i*_ and *E*_*i*_ position, coupling with the environment may have a role in shaping the *S*_*i*_(*E*_*i*_) curves. Therefore, we studied the system at a range of *E*_*e*_ and observed that increments in *E*_*e*_ lead to increased 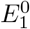 and diminished *γ*_1_ and 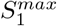, indicating that stronger coupling with the surroundings relaxes the response of the system to changes in *E*_1_ (described below). Thus, the overall phenotypic variation within a given genotype can be decomposed into an internal component arising from stochastic intracellular fluctuations, tissue/season influences, genome variations and metabolic constraints that modify Σ*S*_*i*_, and an external component related with system’s coupling to its surroundings mediated by *E*_*e*_ levels.

## IV. PARAMETERS OF THE LOGISTIC EQUATION

The main characteristics of the logistic curve (Fig. 1) are the asymptotic 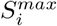 when approaching the highest value of *E*_1_, the inflection point 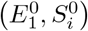, and the sensitivity of change represented by *γ*_*i*_. Therefore, we studied each of these parameters separately in relation with internal (Σ*S*_*i*_) and external (*E*_*e*_) environmental influence.

The asymptotic 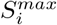 obtained at different *E*_*e*_ levels for each metabolite gave linear relationships for all Σ*S*_*i*_ tested, whose slopes for 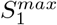 and 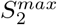 cancel each other (Fig.2A,B). Moreover, the ordinates at *E*_*e*_ = 0, which corresponds to the closed system, are the equilibrium 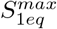 and 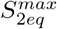. This result prompted us to see if there is a relationship between the degree of displacement from equilibrium [34]

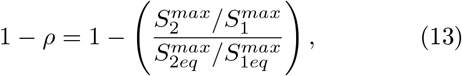

and *J*_*e*_. Interestingly, we obtained a linear relationship that passes through the origin with a positive slope (Fig.2C), suggesting that displacement from equilibrium may be considered a driving force of the metabolic flux. In a sense, we could say that the system operates in a linear regime, despite its overall complexity.

**Figure 2:**
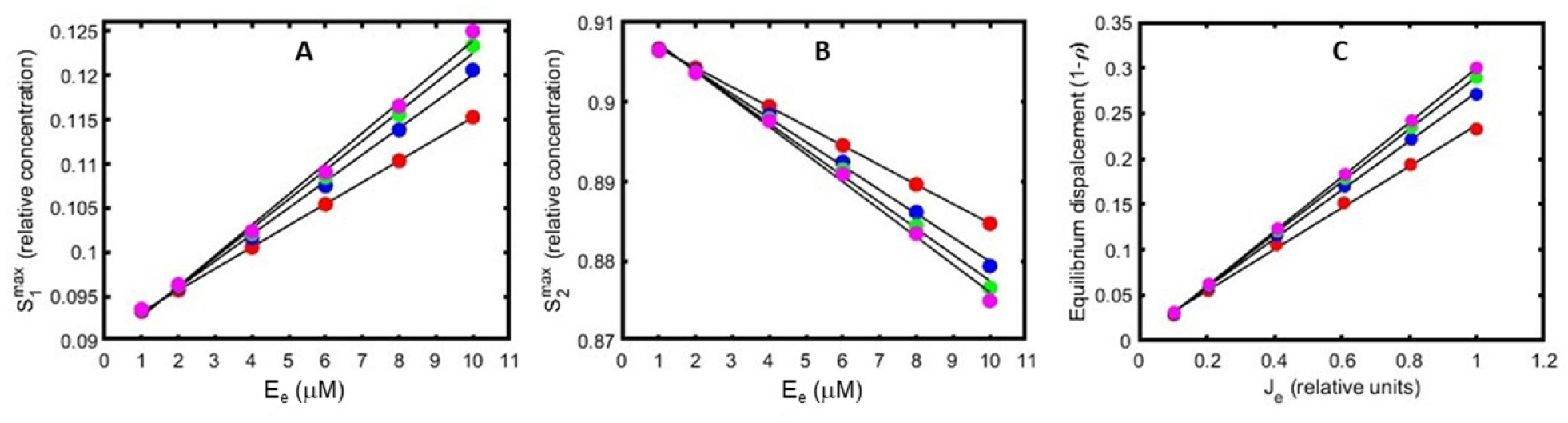
Relationship between 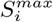 and external flux. A-B: 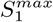 and 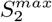 respectively, as function of *E*_*e*_. C: Equilibrium displacement 1 *– ρ* as function of *J*_*e*_. In all graphs, dots are the calculated values with Σ *S*_*i*_ = 0.2 *mM* (red), 2.0 *mM* (blue), 6.0 *mM* (green), and 20.0 *mM* (magenta), and black lines are the adjusted linear relationships, with (0.98 *≤ R*^2^ *≤* 0.99) for A, B, and C. Relative 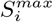 were calculated as in Fig. 1.

The position of the inflection point 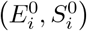 seems to displace as Σ*S*_*i*_ increases (Fig.1, and Supplementary Fig. S3). Hence, we asked if this displacement might be related with reaction order. For this analysis, we considered the position of the inflection point as the vector 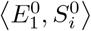. With the idea of associating the displacement to only a scalar, we concentrated on the study of the module of the vector 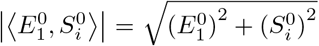. For a range of Σ *S*_*i*_ we observed that *J*_*e*_ responded to increasing 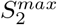 as expected from the irreversible Michaelis-Menten function (Fig.3A). Although 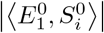 was linearly related to 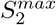 (Fig.3B), its relation with external flux saturation degree, measured as 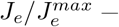 where 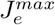 is the equivalent to *V*_*max*_ of the isolated system− was complex. In general, 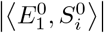 exponentially increased against increases in 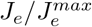, but the parameters that defined the exponential fit changed when the reaction kinetics transitioned from mixed order to zero order (Fig.3C). This trend was more clearly observed when the exponential curve was linearized by taking the natural logarithm of the module of the inflection point (Fig.3D). These results, together with those of Fig.1, indicate that the shape of the *S*_*i*_(*E*_*i*_) logistic curve, and therefore, *d*, depend on the total concentration Σ*S*_*i*_ present in the metabolic pathway, which, in turn, is determined by cell/tissue physiology. This might lead to differences in additive and dominant responses against gene dosage among different tissues.

Studying the derivative of the function (12) and evaluating it at the inflection point 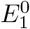, the parameter *γ*_*i*_ may be written as

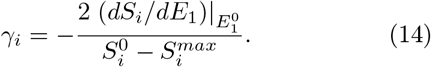

This expression allows us to associate *γ*_*i*_ with the rate of change at the inflection point, characterizing the response of the pathway to enzymatic change. Therefore, the higher the value of *γ*_*i*_, the steeper the curve of *S*_*i*_ against *E*_*i*_ before approaching 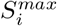values. With this in mind, we wanted to see how *γ*_*i*_ might depend on coupling to the surroundings through *E*_*e*_ values. As may be seen in Fig.4A, *γ*_*i*_ decreases as *E*_*e*_ increases, thus suggesting that, for this simple system, stronger coupling to the surroundings relaxes the metabolite response to changes in internal enzyme levels.

To describe the behavior of *γ*_*i*_ in terms of external enzyme variation, data in Fig. 4A may be fitted with the following linear fractional function

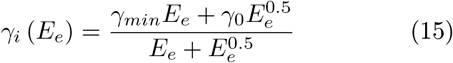

where *γ*_*i*_ (*E*_*e*_ = 0) = *γ*_0_ corresponds to the sensitivity of the isolated system, *γ*_*min*_ is associated to the horizontal asymptote for highest *E*_*e*_ thus representing the asymptotically minimum *γ*_*i*_ value, and *E*^0.5^ symbolizes the external enzyme concentration *E* _*e*_ for which 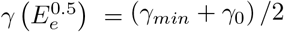. We can see that *γ*_0_, that is, when the system is isolated, increases sharply with Σ*S*_*i*_. Since Σ*S*_*i*_ may be disturbed after a small intake or leak of metabolite, this result indicates that this system is rather unstable when closed and, conversely, its robustness increases as its coupling to the surroundings becomes stronger.

## V. REINTERPRETATION OF ADDITIVE AND DOMINANCE GENE EFFECTS

The maximum amplitude between two *S*_*i*_ values is that between the asymptotic *S*_*i*_ attained at lowest and highest *E*_*i*_, which we will call 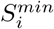 and 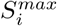 respectively (Supplementary Fig. S1). These *S*_*i*_ values should correspond to extreme *P*_1_ and *P*_2_ contrasting parents (*e*.*g*. see Fig.1 for Σ*S*_*i*_ = 20.0 *mM*). Since our logistic function is symmetrical (Supplementary Fig. S1), and *a*_*i*_ is the half-difference in *S*_*i*_ for the genotypes *g*_*i*_*g*_*i*_ and *G*_*i*_*G*_*i*_, the highest half-difference between two *S*_*i*_ values at any Σ*S*_*i*_ should be 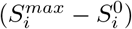, which is the highest *a*_*i*_ value for that Σ*S*_*i*_ (Fig.1). We will call this particular *a*_i_ value as the maximal additive gene effect *A*_*i*_. Hence, we may reinterpret the scaling factor in (14) as −*A*_*i*_ and, therefore, *A*_*i*_ may be expressed as:

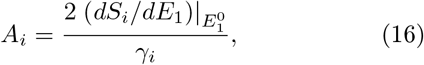

thus underscoring the relationship of additive gene effects with system’s sensitivity. Since, as observed above, *γ*_*i*_ diminishes with increments in *E*_*e*_, a parallel drop in *A*_*i*_ should be observed, as shown in Supplementary Fig. S4, which indicates that *A*_*i*_ diminishes at stronger coupling to the surroundings. Moreover, any *a*_*i*_ value can be divided by *A*_*i*_ to get a fractional number *η*_*i*_ = *a*_*i*_*/A*_*i*_ which may indicate the selectability of the trait, being it 0 ≤ *η*_*i*_ ≤ 1. In addition, this indicates that *η*_*i*_ may depend on Σ*S*_*i*_, and on the curve region where the parents *G*_*i*_*G*_*i*_ and *g*_*i*_*g*_*i*_ lay, i.e., their relative distance from the inflection and asymptotic points.

The relationship between *a*_*i*_ and *d*_*i*_ may be expressed in terms of the ratio of dominance *r*_*i*_, which provides relative value for *d*_*i*_:

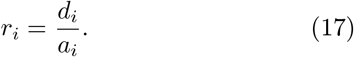

If both parents have *S*_*i*_ values in the concave upward region (see the Fig. 1), *r*_*i*_ will be negative, whereas if *S*_*i*_ values of both parents are in the concave downward region, *r*_*i*_ will be positive, thus being −1 ≤ *r*_*i*_ ≤ 1. Considering that relationship (12) represents a continuous and derivable function, we can establish a connection between *E*_1_ and *r*_2_. In this sense, if we consider that 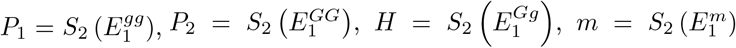 and 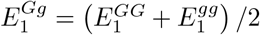, using the Mean Value Theorem [35] we can write

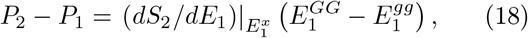

where 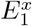 is within the 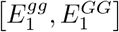 and 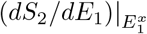 is the derivative of the function *S*_2_ evaluated at 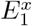. As shown in the Supplementary Material, relationship (18) allows us to find a relationship between *r*_*i*_ and the enzyme concentrations 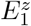 (with *z* = *Gg, gg, m*) given by

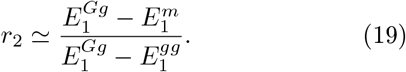

Thinking of equation (19) as an equality, we can calculate the enzyme concentration for which *S*_2_ is at the midpoint between the parents,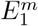:

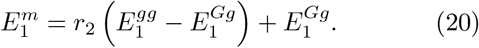

Hence, if dominance is complete, *r*_2_ = 1 and therefore, 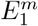 approaches 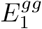. If, on the contrary, there is no dominance, *r*_2_ = 0, and hence 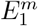 is near 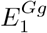. In addition, when 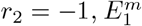 approaches 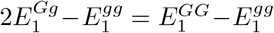 These results support previous findings and highlight that, in this context, whether *d* is complete, partial, negative or null depends on the region of the *S*_*i*_ (*E*_1_) curve where the parent and hybrid genotypes are located relative to the inflection point. It also depends on Σ *S*_*i*_, as it determines the inflection point’s position (Fig. 3B). Nevertheless, this explanation does not rule out the complexity of allelic interactions that ground different cases of dominance [2, 4], and does not include the cases of codominance.

**Figure 3:**
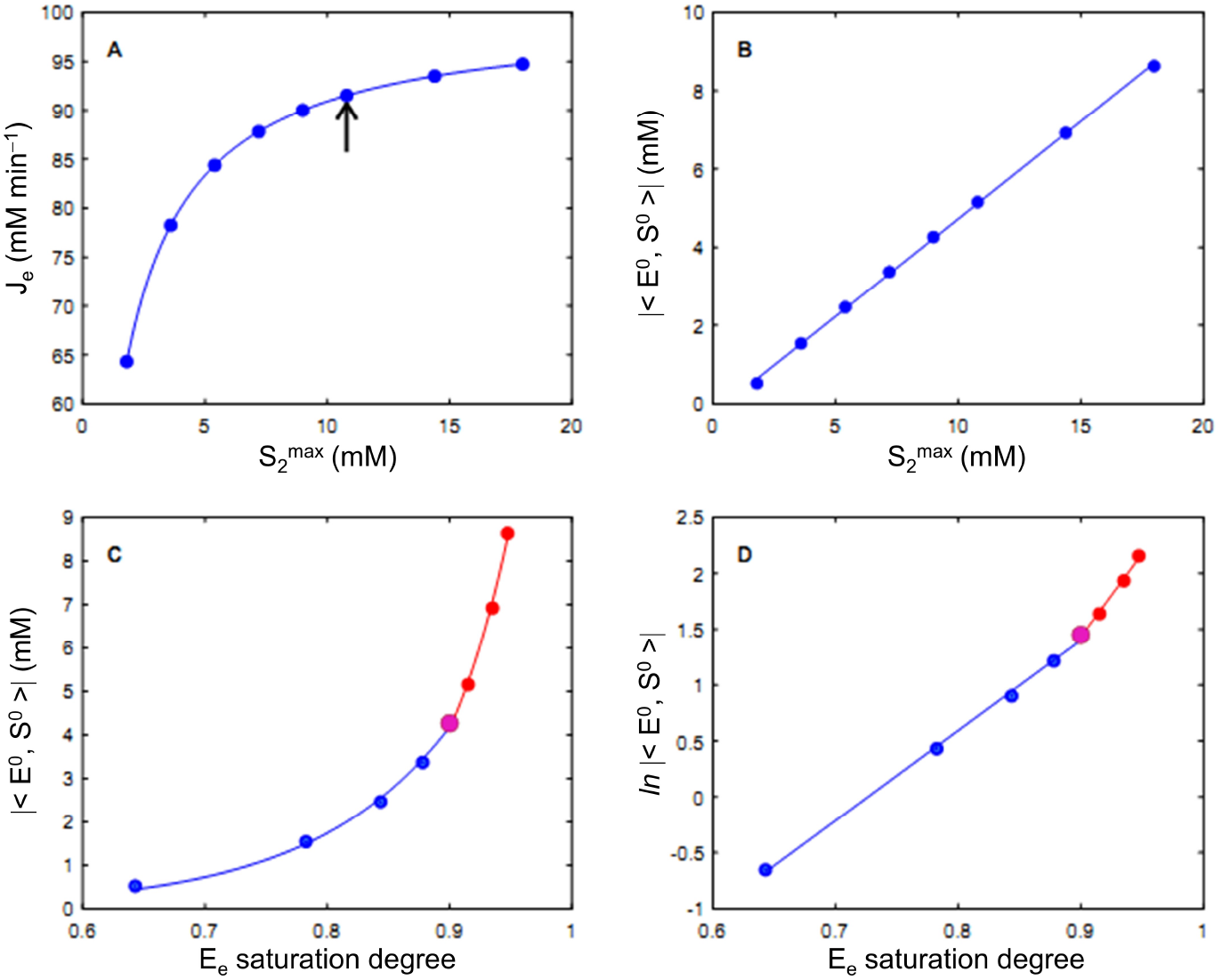
Inflection point displacement as determined by total metabolites concentration. A: Steady-state *J*_*e*_ against 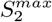 , which follows the irreversible Michaelis-Menten equation, with 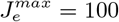 *mMmin*^*−*1^ and *K*_*M*_ = 1.0 *mM*. B: Inflection point module against 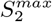, showing a linear relationship. C: Inflection point module against *E*_*e*_ saturation degree 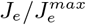, which has an exponential trend with a change in parameters when the kinetics shifts from mixed order (blue) to zero order (red). The violet dot represents the shift, which is indicated with an arrow in A. D: Linearization of data from C, where the shift may be better appreciated. The Σ *S*_*i*_ were 2, 4, 6, 8, 10, 12, 16, and 20 *mM*. The Σ*S*_*i*_ = 0.2 *mM* was not used here because in this case the inflection point has negative coordinates. Dots: calculated values; lines: adjusted reversible Michaelis-Menten equation (A), linear regression (B) quadratic fit (C), and log-transformation (D). In all cases, 0.98 *≤ R*^2^ *≤* 0.99.

**Figure 4:**
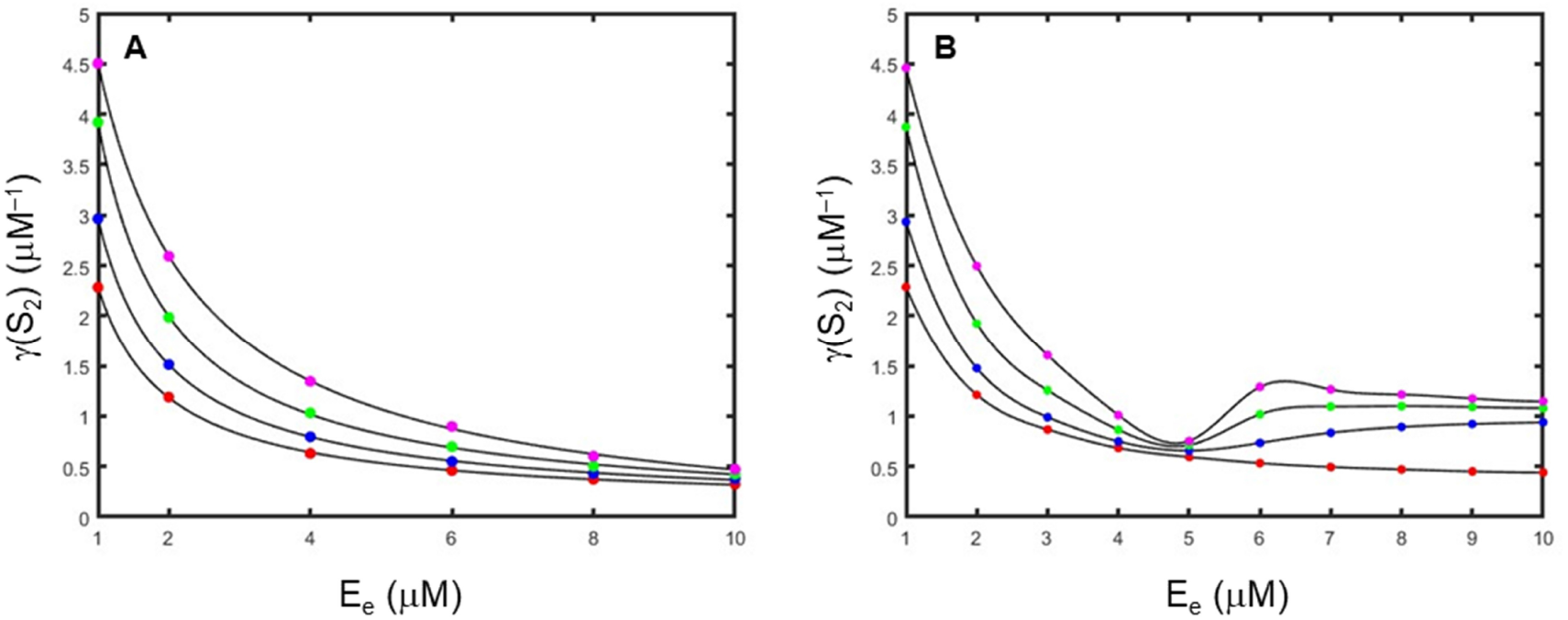
*γ*_*i*_ values as a function of *E*_*e*_ for a pathway with a single step (A) or 4 steps (B). In both, initial conditions were *S*_*i*_ (0) = 0.1 *mM* (red), 1.0 *mM* (blue), 3.0 *mM* (green), and 10.0 *mM* (magenta), givLing rise to Σ *S*_*i*_ = 0.2, 2.0, 6.0, and 20.0 *mM* (A) or 0.5, 5.0, 15.0, and 50.0 *mM* (B) respectively, according to *nS*_*i*_(0) = *ΣS*_*i*_. Dots: *γ*_*i*_ values calculated with eq. 12, black lines: *γ*_*i*_ values adjusted with eq. 15 (0.98 *≤ R*^2^ *≤* 0.99). In B, *E*_*i≠* 4_ = 5 *µM*.

## VI. THE ANALYSIS OF MORE COMPLEX PATHWAYS

To approach more realistic metabolic pathways, we added 1 to 5 reactions to our simple one-reaction path-way (Supplementary Fig. S6). Here, we will focus on a four-reaction pathway. For this analysis we used only Σ *S*_*i*_ = 15 *mM* (*i*.*e. S*_*i*_(0) = 3 *mM*) because for the one-step pathway this condition was in the mixed-order reaction region rendering the logistic curve well defined in the positive quadrant (Figs.1 and 3A), and we assumed that the same properties hold for individual reactions in larger pathways. In a first approach, we varied the concentration of internal enzyme catalyzing the last reaction, *E*_4_, maintaining the concentrations of the other *E*_*i≠* 4_ constant at 5 *µM*.

The last reaction had the same logistic behavior for substrate *S*_4_ and product *S*_5_ against changes in *E*_4_ as previously observed for the one-reaction pathway; however, new trends may be noticed in the previous reactions (Fig. 5A). At the lowest *E*_4_, *S*_3_ and *S*_4_ concentrate almost all the mass of the system. Then, at small *E*_4_ increases, *S*_1_, *S*_2_, and *S*_3_ start to increase following the sigmoid behavior already seen for the one-reaction pathway. However, after reaching a critical *E*_4_ concentration near 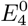, a very sharp peak in *S*_3_ was observed. This may be interpreted taking into account the reversibility of the reactions and the mass conservation of the system. At low *E*_4_, *S*_4_ backward reaction is favored by mass action. Thus, small increments in *E*_4_ lead to higher rates of increment in the concentrations of metabolites at the previous steps than at later steps. Conversely, once *E*_4_ exceeds 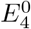, the system shifts toward the forward transformation of *S*_4_ into *S*_5_ and, in this way, further increments in *E*_4_ beyond this point stop accumulation of previous *S*_*i<*4_. These transitions occur abruptly around the inflection point, giving rise to peaks in the concentration profiles of metabolites at previous steps. Consequently, the presence of these peaks is indicative of a regime shift in the system dynamics, centered at the inflection point of *S*_4_ and *S*_5_, which is also manifested in the preceding reaction steps. This behavior suggested a strong mutual correlation among the concentrations of the five metabolites constituting the pathway, as well as their tight coupling with the environmental conditions. It arises primarily from mass conservation constraints, which ensure that any perturbation in the concentration of one metabolite propagates almost instantaneously throughout the entire network.

**Figure 5:**
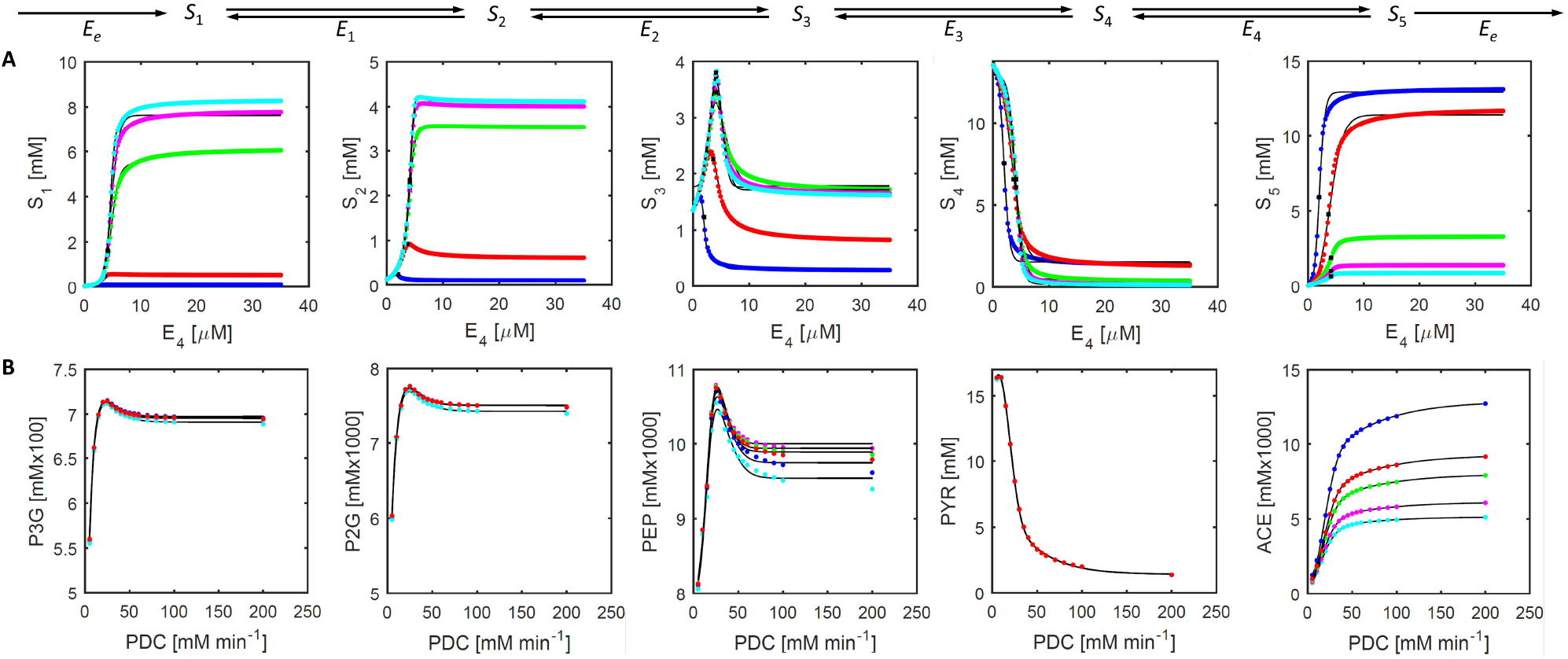
The concentrations of the last five metabolites in a four-reaction pathway schematized on top. A: Fit of *S*_1_-*S*_5_ with eq. 21 when varying *E*_4_. The other enzymes of the pathway, i.e *E*_*i*≠4_, were set to 5 *µM* , with Σ *S*_*i*_ = 15 *mM*. In each graph, *E*_*e*_ = 2 *µM* (blue), 4 *µM* (red), 6 *µM* (green), 8 *µM* (magenta), and 10 *µM* (cyan). The curves adjusted according to eq. (21) are shown as continuous black lines (0.98 *≤ R*^2^ *≤* 0.99). The black asterisk indicates *S*_*i*_ at *E*^0^ (*i*.*e*. the 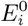 corresponding to the last reaction). B: Metabolites concentrations at the final steps of yeast glycolysis with different pyruvate dehydrogenase (PDC) *V*_*max*_, obtained from gustavsson5 [37] model (dots) and the corresponding curves adjusted with eq. (21) shown in black lines (0.98 *≤ R*^2^ *≤* 0.99). The values of the parameters kLow and kHigh that describe input and output were: 5.17 (blue); 10.17 (red); 15.17 (green); 51.62 (magenta); 51619 (cyan). The default value in gustavsson5 is 516190.

Given that *S*_*i*_(*E*_*i*_) functions are smooth, a straightforward approach to model the peaks observed in Fig. 5A is to augment the logistic function (12) with a Gaussian term:

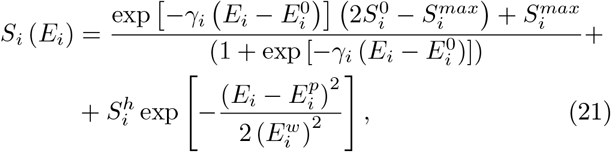

where 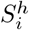 represents the height of the peak on the metabolite axis, 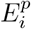 is the enzyme concentration at the peak, and 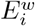 is the width of the peak measured on the enzyme axis. In this manner, the regime change characterized by the emergence of peaks in *S*_*i*_ for *i* = 1, 2, 3 can be effectively captured, seeming the logistic equation (12) a special case that can be obtained when 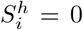. Therefore, equation (21) may be considered a general one for the description of this kind of metabolic pathways. On the other hand, it should be recalled that both 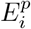 and 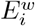 are concentrations of the enzyme catalyzing the last step, in this case, *E*_4_, whereby the effect of this enzyme on the concentrations of metabolites in previous steps is a system effect.

Extending the analysis to other *E*_*i≠*4_, the peak in *S*_3_ was observed in all conditions (Supplementary Fig. S5A); however, the peak widens for *E*_*i≠*4_ = 10 *µM* and *E*_*i≠*4_ = 20 *µM*. In addition, depending on the *E*_*i≠*4_, peaks were observed also in previous reactions. We also noted that the separation of *S*_5_ curves was not even: when 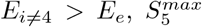 was higher than 10 *mM* thus accounting for most of the mass in the system, but when 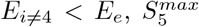 did not surpass half the mass of the system, the other half being shared in the previous reactions; in addition, *S*_3_ peak was attained at 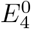 for the cases when *E*_*i≠*4_ *≦ E*_*e*_, while this coincidence occurred at previous reactions when *E*_*i≠*4_ *> E*_*e*_ (compare red and blue curves with the others in Fig. 5A and see also Supplementary Fig. S5A). Furthermore, *γ*_*i*_ decreased as predicted in eq. (15) when *E*_*i≠*4_ *> E*_*e*_ but then, when *E*_*i≠*4_ *< E*_*e*_, *γ*_*i*_ increased, being *E*_*i≠*4_ = *E*_*e*_ the particular enzyme relationship where the switch in the trends of *γ*_*i*_ takes place (Fig. 4B). The change in *γ*_*i*_ trend is apparent when the reactions are occurring with a mixed-order or zero-order kinetics (*i*.*e*. with Σ*S*_*i*_ = 5; 15 or 50 *mM*) but not when the system is at first-order kinetics (Σ*S*_*i*_ = 0.5 *mM*). Taken together, these results indicate the broad consequences of the change of regime arisen at inflection point, and how it is conditioned by Σ*S*_*i*_ and *E*_*e*_.

Furthermore, we analyzed the same pathway with different numbers of steps. As shown in Supplementary Fig. S6, the trend observed for the last five metabolites was exactly matched in all the pathway lengths, but at the beginning of the pathways with 5 or 6 steps the relationship *S*_*i*_(*E*_*i*_) was logistic again, indicating that the change of regime does not propagate backward beyond 3 reaction steps. Moreover, for the case of *S*_*i*_ being the substrate of the third reaction before *E*_*n*_, 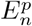 coincides with 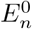, and later, 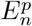 tends to diminish linearly as the step into consideration approaches the catalyzed reaction (Supplementary Fig. S7), indicating that the change of regime at the inflection point is sensed at previous steps.

In view of these results, we wondered about varying other enzymes of the pathway. In Supplementary Fig. S8 we show the four-step pathway where we changed *E*_1_, *E*_2_ or *E*_3_, maintaining the other enzyme concentrations constant, at 20 *µM*. We observed that, as in Fig.5A, the substrate and product of the reaction affected changed with a logistic function and the previous metabolites exhibited the behavior described by eq. (21). However, the reactions downstream of that affected by the change in enzyme concentration continued displaying the logistic behavior only, indicating that the change in regime does not propagate forward.

Finally, it should be taken into account that *T* is not necessarily determined by the substrate or product of the enzyme affected by the genotypic change (in this example, *E*_4_), but it might be also determined by another metabolite of the same pathway. Hence, if *T* is determined by the concentration of a metabolite from a step previous to that catalyzed by *E*_4_, overdominance (*d*_*i*_ *> a*_*i*_) may be observed when enzyme concentrations from the contrasting parents, 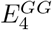 and 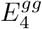, are at opposite sides of 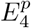 (Supplementary Fig. S9A). On the other hand, dominance reversals may be possible as consequences of changes in *E*_*e*_. For instance, if *T* is determined by *S*_3_ and the parents possess *E*_4_ levels lower than 8 *µM* , it is possible that, for a same pair of 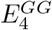 and 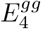, the *G*_4_*G*_4_ parent has higher *T* value than *g*_4_*g*_4_ when *E*_*e*_ = 10 *µM* but lower if *E*_*e*_ = 6 *µM* (Supplementary Fig. S9B), thus indicating that dominance reversal might arise after a change in the connection with the surroundings. This is in agreement with observations that dominance reversals are strongly dependent on the context [36].

## VII. VALIDATION OF THE MODEL

To see if the logistic explanation of additive and dominance gene effects may fit in real situations, we addressed two approaches. On the one hand, we generated data with an independent model based on experimentally determined parameters and kinetic equations to see if these data may be adjusted with equation (21), and on the other hand, we applied equation (12) to fit experimental data obtained from the literature. Following we describe the results obtained.

### A. Adjustment of data generated with an independent model

In this approach, we used the model developed by Gustavsson et al. [37], which may be accessed at JWS Online (https://jjj.mib.ac.uk/models/gustavsson5/simulate/, last visited on 10/15/2026). This model was built in the frame of *silicon cell* algorithms using experimentally validated kinetic equations and parameters [37, 38] to understand the role of phosphofructokinase allosteric control in determining glycolytic oscillations in single yeast cells. We choose the glycolytic pathway because it is well understood and characterized, and allowed us to see if the general trends discussed above may be observed in the last steps of this simulation. To mimic the different *E*_*e*_ levels, we modified the values of the external parameters kLow and kHigh [37] and choose pyruvate dehydrogenase (PDC) as the varying *E*_*i*_, which catalyzes the oxidative decarboxylation of pyruvate (PYR) to render acetate (ACE) and CO_2_. This reaction may be accounted for as one of the final steps in glycolysis, being the last reaction in a linear chain comprising 4 reaction steps. The results obtained for a range of PDC concentrations (expressed as maximum velocity, *V*_*max*_) are shown in Fig. 5B. All data points were satisfactorily fitted by eq. (21). Moreover, we noticed clear similarities with features found in our calculations, which could not have been anticipated before, such as the presence of an inflection point, the diminishing of the product concentration curves at increasing exit fluxes, the superposition of substrate concentration curves and simultaneous separation of product concentration curves at different exit fluxes, and the presence of peaks at PDC concentrations near 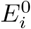 for PEP, P2G and P3G suggesting a change of regime. However, to obtain these curves we had to work at a range of kLow and kHigh below the default value of the gustavsson5 model [37]; otherwise, we observed little variation in ACE concentration as may be inferred from Fig. 5A at *E*_*i≠*4_ *< E*_*e*_ (cyan and magenta curves). We hypothesized that this high exit rates in gustavsson5 default or in the mentioned condition in Fig. 5A was too high to retain the increasing ACE within the cell despite PDC concentration increments, and we corroborated it by running our model with *E*_*e*_ = 100 *µM* (not shown). Taken together, these results indicate that eq. ( 21) is able to fit the changes in metabolite concentrations of the final steps of the gustavsson5 silicon cell simulation (Fig. 5B), which involves reactions governed by diverse experimentally validated kinetic functions and parameters, yet it was successfully described by our simpler model.

### B. Adjustment of observational data

To see if eq. (12) may fit experimental data, we looked for examples complying the following conditions, which are essential for a faithful comparison with our simplified model: 1) that one gene determines a trait in a plant or animal, 2) that this trait is determined by the concentration of a metabolite, 3) that this metabolite is the product of the activity of an enzyme encoded by the gene found in 1), 4) that the said enzyme has Michaelis-Menten kinetics, 5) that this metabolite is produced in a linear pathway, 6) that this metabolite is the last in the pathway and that the trait in question depends on how much of this metabolite reaches a specific sink, 7) that there exist paired data for enzyme activity and metabolite concentration, and 8) that there are at least five paired data points available of the type mentioned in 6). After reviewing the literature, we found as only example the synthesis of amylose in plant tissues catalyzed by granule-bound starch synthase I (GBSSI). Amylose is part of starch, one of the most important energy reserve compounds in photosynthetic organisms, and essential for human nutrition from plant tissues such as tubers, fruits, seeds and their products. In starch granules, amylose is present alongside amylopectin, primarily in the amorphous regions. It is the final product of a pathway that seems linear in its last steps, which comprise the conversion of glucose-1-phosphate into ADP-glucose catalyzed by ADP-glucose pyrophosphorylase followed by incorporation of the glucose moiety by glycosidic *α* − 1, 4 linkages into the growing amylose chain, catalyzed, in starch granules, by GBSSI [39]. This enzyme has a typical Michaelis-Menten kinetics as that used in our model [40], so all these properties make this pathway the choice for our objective. In this pathway, amylose is incorporated accompanying starch granule growth, and for this reason amylose contents (AC) are determined in starch granules and commonly expressed as percentage of starch.

Since eq. (12) has 4 parameters, any attempt to adjust real data needs datasets with at least 5 AC-GBSSI paired concentrations and, in addition, some AC values should approach asymptote for anchoring one of the parameters. Looking at the literature, we found several examples with such datasets where, in general, there was a trend of increasing AC with increasing GBSSI activity. However, since experiments were not designed in the past taking into account eq. (12), some reports present too dispersed data, others choose examples with low, middle or high AC that cannot be adjusted because there are less data points than parameters, and others looked for polynomial adjustments between GBSSI and AC (*e*.*g*. ref. [41, 42]).

In Fig. 6 we present examples in six different plant species where at least five data points were determined, and where data approaching AC asymptotic value was seeming, which is not an unreasonable assumption given that the cell compartment where the reaction takes place−in this case the starch granule−has a limited volume. Riewe et al. [43] manipulated the level of apyrase in potato tubers to investigate the metabolic adaptations to decreasing ATP contents. As part of the responses, they found that GBSSI changed in 5 transgenic potato lines overexpressing an *Escherichia coli* apyrase gene and, in response, AC changed as predicted by eq. (12) (Fig. 6A). In Arabidopsis, a study of natural polymorphisms of GBSSI [42] also found increases in AC in relation with increases in GBSSI activity. However, the changes in GBSSI activity were due both to differences in enzyme contents as well as to differences in GBSSI specific activity measured as units of activity per mass unit of protein with purified enzyme *in vitro*. Although all data were also fit by eq. 12, we represented in Fig. 6B only those 6 genotypes in which purified GBSSI specific activity was similar, in such a way that the differences in total activity per mass of plant tissue were due to differences in enzyme contents between genotypes. Conversely, in rice, Liu et al. [44] produced aminoacids substitutions in GBSSI by site-directed mutagenesis to obtain allels with a range of specific activities, while maintaining the same amount of total protein in all the mutant lines. In Fig. 6C the adjustment of their data of GBSSI and AC from seven lines with eq. (12) is shown. In banana fruits GBSSI and AC were quantified in a single variety during fruit development and posthaverst [45]. The changes in GBSSI activity along these periods were reflected in AC, fitting well with eq. (12) (Fig. 6D). In millet, Hunt et al. [46] studied the GBSSI activity in the S and L *loci* by analyzing 6 genotypes and their data fit again with eq. (12) (Fig. 6E). In wheat, Zi et al. [47] induced mutations in GBSSI by treatment with ethyl methane sulfonate (EMS) and quantified GBSSI and AC in seven mutants and the wild type, which again show a very good fit with eq. (12) (Fig. 6F).

**Figure 6:**
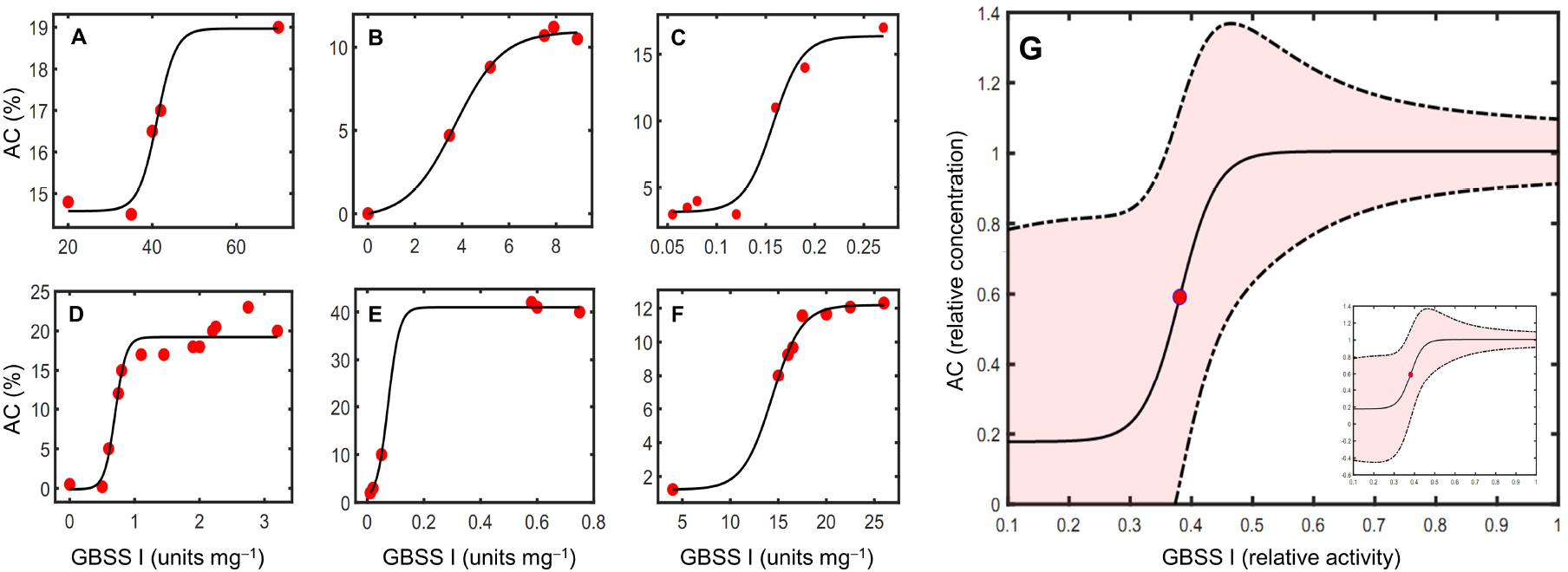
Experimentally determined amylose contents (AC) at different granule-bound starch synthase I (GBSSI) specific activities per unit of tissue mass in plants (0.98 *≤ R*^2^ *≤* 0.99). A: potato [43]; B: Arabidopsis [42]; C: rice [44]; D: banana [45]; E: millet [46] ; F: wheat [47]. Red dots are experimental values obtained from the indicated references, and black lines are fits obtained with eq. (12). G: curves obtained using eq. (12) with the average parameters from relative values (continuous line) *±*95% confidence interval (dotted lines). The red dot indicates the inflection point of the mean curve. In the *inset*, the negative *S* values at the low limit of confidence interval*−*which have no physical sense*−*are shown. Pink-shadowed area indicates expected relative (*S, E*) mean coordinates in plants within this confidence interval.

From these curves we calculated the parameters *γ, E*^0^, *S*^0^, and *S*^*max*^ for GBSSI and AC in each plant, as well as their mean ± standard deviation (SD) in an attempt to obtain a more general picture of enzyme-product relationships in this pathway (Table I).

**Table I:** Parameters of logistic curves from Fig. 6 for AC in six plant species.

| Parameter | P | A | R | B | M | W | Mean ± SD | CV |
| --- | --- | --- | --- | --- | --- | --- | --- | --- |
| $E^0$ | 41.14 | 3.70 | 0.16 | 0.70 | 0.07 | 14.35 | 10.02±16.19 | 161.55 |
| $S^0$ | 16.77 | 5.34 | 9.76 | 9.55 | 20.67 | 6.71 | 11.47±5.99 | 52.27 |
| $S^{max}$ | 18.97 | 10.95 | 16.46 | 19.22 | 41.00 | 12.19 | 19.80±10.93 | 55.23 |
| $\gamma$ | 0.45 | 0.99 | 67.43 | 12.09 | 49.80 | 0.66 | 21.90±29.31 | 133.84 |
| $A$ | 2.20 | 5.61 | 6.70 | 9.68 | 20.33 | 5.48 | 8.33±6.36 | 76.20 |

P: potato; A: Arabidopsis; R: rice; B: banana; M: millet; W: wheat. CV: coefficient of variation, SD/Mean (%)

We noticed that, while SD of AC-related parameters *S*^0^ and *S*^*max*^ was around 50-60% of mean, for the GBSSI-related parameter *E*^0^, SD surpassed 160%. As a consequence, *γ*, which is also related to GBSSI, had SD around 130% of its mean. This high dispersion in GBSSI-related parameters may be explained not only by the diversity of plant species, but mostly because of the different conditions under which GBSSI activity values were obtained −including source tissue, plant age, methods for extraction and activity measuring, etc. By contrast, AC measures are less prompt to dispersion because all them were obtained from starch granules with a more uniform methodology. Despite this dispersion, it may be appreciated in Fig. 6A-F that the trends were similar. Hence, to get a better comparison between plants we calculated the relative values of GBSSI activity and AC by dividing each individual value by the highest value recorded for the correspondig plant, which we used to calculate the mean parameters and curve (Fig. 6G). Furthermore, from each plant’s curve we constructed a matrix of covariance that correlates the parameter values of each individual curve, from which we generated random curves using Cholesky decomposition [48]. We also used a filtering procedure to discard those random curves for which *S*^*max*^ ≤ *S*^0^ or *γ* ≤ 0. After obtaining 50,000 random curves in this way, we calculated SD and confidence interval (95%), which is represented, together with the mean curve, in Fig. 6G. In addition, we calculated mean *A* as 8.3 ± 6.0 (95% confidence interval, 4 *df*) in this system, indicating that mean increases/decreases in AC by selection may be expected within this interval. Taken together, these data show that eq. (12) may represent the variation of AC against changes in GBSSI activity in plants provided datasets comply with the requisites indicated above. More generally, we conjecture that this behavior holds true whenever the metabolic pathway in question can be considered linear with a saturable coupling to its environment.

## VIII. CONCLUDING REMARKS

In this study, we modeled a simplified metabolic pathway consisting of monosubstrate reactions catalyzed by monomeric enzymes that obey Michaelis–Menten kinetics through a system of coupled ordinary DEs. Despite its simplifying assumptions, this framework is supported by independent *in silico* simulations and experimental observations. This is particularly relevant because the above simple assumptions did not preclude the interpretation of metabolic systems characterized by more complex enzyme kinetics and quaternary structures. Furthermore, the reduced complexity of the model enabled us to identify key parameters and trends.

The stationary state out of equilibrium was achieved analytically by imposing three conditions: 1) opening the system to entrance and exit of matter catalyzed by saturating enzyme kinetics, 2) exactly matching the input and output rates, and 3) limiting the total mass of the system, Σ*S*_*i*_. As a result, the system was described by a logistic *S*_*i*_(*E*_*i*_) relationship with four parameters: the asymptotic metabolite concentration 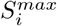, the system sensitivity to enzymatic changes *γ*_*i*_, and the metabolite and enzyme concentrations at the inflection point, 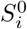 and 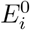 respectively. In particular, *γ*_*i*_ depends both on Σ *S*_*i*_ , which characterizes the internal state of the system, and on the external flux *J*_*e*_, which reflects the strength of the system’s coupling to its environment. Moreover, in a chain of reactions with more than two metabolites, we noticed a change of regime in the reactions preceding that one where *E*_*i*_ varies, which may be observed in the shape of the curves and in the trend of *γ*_*i*_. This change of regime was modeled by inserting a Gaussian function with three additional parameters that fully describe all the catalyzed steps of the pathway. A reduction in the number of these parameters was not possible because 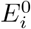 and 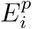 were not always coincident. Of particular relevance is the observation that the logistic relationship holds regardless of the number of intermediates in the pathway and also at different strengths of pathway coupling with its surrounding environment.

We further validated the framework by fitting it to data from an independent silicon cell model and from experimental observations in plants. The quality of these fits suggests that whenever at least five paired *S*_*i*_ and *E*_*i*_ values can be experimentally determined−including some *S*_*i*_ values approaching saturation−the parameters 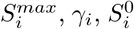, and 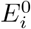 may be estimated without requiring detailed knowledge of the kinetic models or parameters underlying each enzymatic reaction. Thus, a macroscopic description of how the metabolic pathway underlying a trait of interest determines additive and dominance gene effects may be achieved more readily.

Our framework provides a reinterpretation of additive and dominance gene effects in relation to morphological traits. These concepts were originally developed from a statistical perspective, at a time when their molecular underpinnings were unknown. However, now it is clear that, while additivity is deterministic, dominance is an emergent property from the complex metabolic and regulatory networks. Consequently, they are logically and functionally heterogeneous (*i*.*e*., they operate at different levels of organization). Regarding additivity, we defined the maximal additivity *A*_*i*_ as the difference 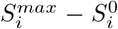, which may be regarded as a system parameter describing the relationship between the asymptotic values and the inflection point. The response to artificial selection depends strongly on additive gene effects because dominance is not directly inheritable. Accordingly, the additive value *a*_*i*_, which depends on the (*S*_*i*_, *E*_*i*_) of the contrasting parents *G*_*i*_*G*_*i*_ and *g*_*i*_*g*_*i*_, may be related to *A*_*i*_ for the given *S*_*i*_. Thus, the ratio *η*_*i*_ = *a*_*i*_*/A*_*i*_, which we call selectability, quantifies how close the observed additive effect is to its theoretical maximum and, consequently, how much scope remains for further selection. Since the (*S*_*i*_, *E*_*i*_) coordinates themselves depend on Σ*S*_*i*_, *E*_*e*_, and *γ*_*i*_, these parameters ultimately determine the selectability *η*_*i*_ of trait *T*.

In the case of dominance, within our limited framework we were able to offer only a correlation of the different cases of dominance with the shape of the *S*_*i*_(*E*_*i*_) region of the curves where the contrasting parents and the hybrid are located, without pretending to rule out other explanations based on system properties and the vast effects of epistasis, pleiotropy and genetic background on phenotypic value [4–6]. Nevertheless, different degrees of dominance may be found just postulating changes in Σ*S*_*i*_ (Fig.1) and, in addition, depending on the connection with the surroundings, dominance reversals and overdominance may also be explained if *T* is caused by an *S*_*i≠n*_ (Fig. 5 and Supplementary Fig. S9). Therefore, the nuance introduced by considering the logistic *S*_*i*_(*E*_*i*_) relationship in the analysis of dominance may facilitate phenotypic analysis if one assumes that phenotype is, ultimately, the result of metabolite concentrations (in the broad sense adopted here) in a given compartment. This explanatory power resides in taking into account the decisive influence of changes in total metabolite concentration Σ*S*_*i*_−which is often overlooked−in addition to changes in *E*_*i*_. Moreover, our framework also considers the role of coupling to the environment by making *J*_*e*_ as depending on the concentration of the last metabolite of the pathway, showing that internal parameters and, as consequence, additive and dominance relationships, also depend on *J*_*e*_. Therefore, despite its simplicity, the present model indicates that changes in enzyme concentration in response to gene dosage are not the only driver of dominance.

Although our model deliberately excludes regulatory mechanisms [49], it can be viewed as an extension of the classical Kacser and Burns [9] framework. Importantly, gene expression —encompassing RNA synthesis, degradation, aminoacyl-tRNA synthesis, polypeptide polymerization, and protein inactivation and degradation— may be considered as the result of complex enzyme-catalyzed metabolic pathways [50]. In addition, regulation of gene expression operates not as a simple on–off switch, but rather through modulation by association constants among macromolecules (proteins, RNAs, and DNA). Thus, the models of Omholt et al. and Kacser and Burns are not mutually exclusive; rather, they may be unified within a broader framework that incorporates both metabolic and regulatory layers in a multiplex network constructed using the principles described here. This unified perspective offers a promising avenue for future work. Subsequent studies will determine whether this framework may prove useful in practical applications such as animal and plant breeding.

## Supporting information

Analytical foundations and Figs. S1-S9

## Acknowledgments

AM and ARL are full professors at Universidad Nacional de La Plata, Argentina. ARR is head of practical assignments at Universidad Nacional de La Plata, Argentina. ARL is principal researcher at CONICET, Argentina.

## References

[1] Burch J, Chin M, Fontenot BE, Mandal S, McKnight TD, Demuth JP, Blackmon H. (2024) Wright was right: leveraging old data and new methods to illustrate the critical role of epistasis in genetics and evolution. Evolution 78:624–634. doi:10.1093/evolut/qpae003.

[2] Di C, Lohmueller KE. (2024) Revisiting dominance in population genetics. Genome Biol Evol 16:evae147. doi:10.1093/gbe/evae147.

[3] Kearsey MJ, Pooni HS. (1996) The Genetical Analysis of Quantitative Traits. Chapman and Hall, London. doi:10.1007/978-1-4899-4441-2).

[4] Veitia RA, Caburet S, Birchler JA. (2018) Mechanisms of Mendelian dominance. Clin Genet 93:419–428. doi:10.1111/cge.13107.

[5] Zschocke J, Byers PH, Wilkie AOM. (2023) Mendelian inheritance revisited: dominance and recessiveness in medical genetics, Nat Rev Genet 24:442–463. doi:10.1038/s41576-023-00574-0.

[6] Tautz D, Pallares LF, Andersson L, Barghi N, Barton N, Bay R, Chan YF, Hancock A, Kaiser TS, Koenig D, Kontarakis Z, Liedvogel M, de Meaux J, Nordborg M, Palmer AA, Purugganan M, Schlötterer C, Schmid K, Stainier DYR, Weigel D, Wolf JBW, Ebert D, Gibson G. (2026) Beyond Mendel: a call to revisit the genotype-phenotype map through new experimental paradigms. Genetics 232:iyag024. doi:10.1093/genetics/iyag024.

[7] Sussmilch FC, Ross JJ, Reid JB. (2022) Mendel: From genes to genome Plant Physiol 190:2103–2114. doi:10.1093/plphys/kiac424.

[8] Feng, C., Chen, B., Hofer J. Shi Y, Jiang M, Song B, Cheng H, Lu L, Wang L, Howard A, Bendahmane A, Fouchal A, Moreau C, Sawada C, LeSignor C, Zhang C, Vikeli E, Tsanakas G, Zhao H, Cheema J, Barclay JE, Hou J, Sayers L, Wingen L, Vigouroux M, Vickers M, Ambrose M, Dalmais M, Higuera-Poveda P, Li P, Yuan Q, Spanner R, Horler R, Wouters R, Chundakkad S, Wu T, Zhao X, Li X, Sun Y, Huang Z, Wu Z, Deng XW, Steuernagel B, Domoney C, Ellis N, Chayut N, Cheng S. (2025) Genomic and genetic insights into Mendel’s pea genes. Nature 642:980–989. doi:10.1038/s41586-025-08891-6.

[9] Kacser H, Burns JA. (1981) The molecular basis of dominance. Genetics 97:639–666. doi:10.1093/genetics/97.3-4.639.

[10] Agutter PS. (2008) The flux-summation theorem and the “evolution of dominance”. J Theor Biol 254:821–825. doi:10.1016/j.jtbi.2008.07.027.

[11] de Vienne D, Coton C, Dillmann C. (2023) The genotype-phenotype relationship and evolutionary genetics in the light of the Metabolic Control Analysis. Biosystems 232:105000. doi:10.1016/j.biosystems.2023.105000.

[12] Raamsdonk LM, Teusink B, Broadhurst D, Zhang N, Hayes A, Walsh MC, Berden JA, Brindle KM, Kell DB, Rowland JJ, Westerhoff HV, van Dam K, Oliver SG. (2001) A functional genomics strategy that uses metabolome data to reveal the phenotype of silent mutations. Nat Biotechnol 19:45–50. doi:10.1038/83496.

[13] Cornish-Bowden A, Cárdenas ML. (2005) Systems biology may work when we learn to understand the parts in terms of the whole. Biochem Soc Trans 33:516–519. doi:10.1042/BST0330516.

[14] Reddy ASN, Huang J, Syed NH, Ben-Hur A, Dong S, Gu L. (2020) Decoding co-/post-transcriptional complexities of plant transcriptomes and epitranscriptome using next-generation sequencing technologies. Biochem Soc Trans 48:2399–2414. doi:10.1042/BST20190492.

[15] Pezoulas VC, Hazapis O, Lagopati N, Exarchos TP, Goules AV, Tzioufas AG, Fotiadis DI, Stratis IG, Yannacopoulos AN, Gorgoulis VG. (2021) Machine learning approaches on high throughput NGS data to unveil mechanisms of function in biology and disease. Cancer Genom Proteom 18:605–626. doi:10.21873/cgp.20284.

[16] Borah K, Das HS, Seth S, Mallick K, Rahaman Z, Mallik S. (2024) A review on advancements in feature selection and feature extraction for high-dimensional NGS data analysis. Funct Integr Genomics 24:139. doi:10.1007/s10142-024-01415-x.

[17] Kemble H, Nghe P, Tenaillon O. (2019) Recent insights into the genotype-phenotype relationship from massively parallel genetic assays. Evol Appl 12:1721–1742. doi:10.1111/eva.12846.

[18] Markin CJ, Mokhtari DA, Sunden F, Appel MJ, Akiva E, Longwell SA, Sabatti C, Herschlag D, Fordyce PM. (2021) Revealing enzyme functional architecture via high-throughput microfluidic enzyme kinetics. Science 373:eabf8761. doi:10.1126/science.abf8761.

[19] Wei H, Li X. (2023) Deep mutational scanning: A versatile tool in systematically mapping genotypes to phenotypes. Front Genet 14:1087267. doi:10.3389/fgene.2023.1087267.

[20] Jain A, Teshima M, Buryska T, Romeis D, Haslbeck M, Döring M, Sieber V, Stavrakis S, de Mello A. (2024) High-throughput absorbance-activated droplet sorting for engineering aldehyde dehydrogenases. Angew Chem Int Ed Engl 63:e202409610. doi:10.1002/anie.202409610

[21] Bost B, Dillmann C, de Vienne D. (1999) Fluxes and metabolic pools as model traits for quantitative genetics. I. The L-shaped distribution of gene effects. Genetics 153:2001–2012. doi:10.1093/genetics/153.4.2001.

[22] Park JO, Rubin SA, Xu YF, Amador-Noguez D, Fan J, Shlomi T, Rabinowitz JD. (2016) Metabolite concentrations, fluxes and free energies imply efficient enzyme usage. Nat Chem Biol 12:482–489. doi:10.1038/nchem-bio.2077.

[23] Dourado H, Mori M, Hwa T, Lercher MJ (2021) On the optimality of the enzyme–substrate relationship in bacteria. PLoS Biol 19:e3001416. doi:10.1371/journal.pbio.3001416.

[24] Cornish-Bowden A. (2012) Fundamentals of Enzyme Kinetics. Wiley-VHC Verlag & Co. Weinheim. ISBN: 978-3-527-33074-4.

[25] Murray JD. (2002) Mathematical Biology: I. An Introduction. 3rd ed. Springer-Verlag New York. doi:10.1007/b98868.

[26] Friedman N, Cai L, Xie XS. (2006) Linking stochastic dynamics to population distribution: An analytical framework of gene expression”, Phys Rev Lett 97:168302. doi:10.1103/PhysRevLett.97.168302.

[27] Taniguchi Y, Choi PJ, Li G-W, Chen H, Babu M, Hearn J, Emili A, Xie XS. (2010) Quantifying E. coli proteome and transcriptome with single-molecule sensitivity in single cells. Science 329:533–538. doi:10.1126/science.1188308.

[28] Chari S, Dworkin I. (2013) The conditional nature of genetic interactions: The consequences of wild-type backgrounds on mutational interactions in a genome-wide modifier screen. PLoS Genet 9:e1003661. doi:10.1371/journal.pgen.1003661.

[29] Fukutomi Y, Phillips-García A, Liu J, Chuang A, Watada M, Ramniwas S, Kopp A. (2026) Evolution of dominance in a Mendelian trait is linked to the evolution of environmental plasticity. Evolution 80:108–126. doi:10.1093/evolut/qpaf185.

[30] Gaines TA, Duke SO, Morran S, Rigon CAG, Tranel PJ, Küpper A, Dayan FE. (2020) Mechanisms of evolved herbicide resistance. J Biol Chem 295:10307–10330. doi:10.1074/jbc.REV120.013572.

[31] Percival CJ, Marangoni P, Tapaltsyan V, Klein O, Hallgrímsson B. (2017) The interaction of genetic background and mutational effects in regulation of mouse craniofacial shape. G3 (Bethesda) 7:1439–1450. doi:10.1534/g3.117.040659.

[32] Rosenblum EB, Rompler H, Schoneberg T, Hoekstra H. (2010) The molecular and functional basis of phenotypic convergence. Proc Natl Acad Sci USA 107:2113–2117. doi:10.1073/pnas.0911042107.

[33] San-Jose LM, Ducret V, Ducrest AL, Simon C, Roulin A. (2017) Beyond mean allelic effects: A locus at the major color gene MC1R associates also with differing levels of phenotypic and genetic (co)variance for coloration in barn owls. Evolution 71:2469–2483. doi:10.1111/evo.13343.

[34] Fell D. (1997) Undesrtanding the Control of Metabolism. Portland Press. London. ISBN: 185578047X.

[35] Thomas GB, Finney RL. (1996) Calculus and Analytic Geometry, 9th ed. Adison-Wesley Publishing Company. Reading, Mass. ISBN-10.0201531747.

[36] Grieshop K, Ho EKH, Kasimatis KR. (2024) Dominance reversals: the resolution of genetic conflict and maintenance of genetic variation. Proc. R. Soc. B 291:20232816. doi:10.1098/rspb.2023.2816).

[37] Gustavsson AK, van Niekerk DD, Adiels CB, Kooi B, Goksör M, Snoep JL. (2014) Allosteric regulation of phosphofructokinase controls the emergence of glycolytic oscillations in isolated yeast cells. FEBS J 281:2784–2793. doi:10.1111/febs.12820.

[38] Teusink B, Passarge J, Reijenga KA, Esgalhado E, Van der Weijden CC, Schepper M, Walsh MC, Bakker BM, Van Dam K, Westerhoff HV, Snoep JL. (2000) Can yeast glycolysis be understood in terms of in vitro kinetics of the constituent enzymes? Testing biochemistry. Eur J Biochem 267:5313–5329. doi:10.1046/j.1432-1327.2000.01527.x.

[39] Goren A, Ashlock D, Tetlow IJ. (2018) Starch formation inside plastids of higher plants. Protoplasma 255:1855–1876. doi:10.1007/s00709-018-1259-4.

[40] Sharma V, Jahan K, Kumar P, Puri A, Sharma VK, Mishra A, Bharatam PV, Sharma D, Rishi V, Roy J. (2022) Mechanistic insights into granule-bound starch synthase I (GBSSI.L539P) allele in high amylose starch biosynthesis in wheat (Triticum aestivum L.). Funct Integr Genomics 23:20. doi:10.1007/s10142-022-00923-y.

[41] Itoh K, Ozaki H, Okada K, Hori H, Takeda Y, Mitsui T. (2003) Introduction of Wx transgene into rice wx mutants leads to both high- and low-amylose rice. Plant Cell Physiol 44:473–480. doi:10.1093/pcp/pcg068.

[42] Seung D, Echevarría-Poza A, Steuernagel B, Smith AM. (2019) Natural polymorphisms in Arabidopsis result in wide variation or loss of the amylose component of starch. Plant Physiol 182:870–881. doi:10.1104/pp.19.01062.

[43] Riewe D, Grosman L, Zauber H, Wucke C, Fernie AR, Geigenberger P. (2008) Metabolic and developmental adaptations of growing potato tubers in response to specific manipulations of the adenylate energy status. Plant Physiol 146:1579–1598. doi:10.1104/pp.108.115758.

[44] Liu D, Wang W, Cai X. (2014) Modulation of amylose content by structure-based modification of OsGBSS1 activity in rice (Oryza sativa L.). Plant Biotechnol J 12:1297–1307. doi:10.1111/pbi.12228.

[45] Miao H, Sun P, Liu W, Xu B, Jin Z. (2014) Identification of genes encoding granule-bound starch synthase involved in amylose metabolism in banana fruit. PLoS One 9:e88077. doi:10.1371/journal.pone.0088077.

[46] Hunt HV, Moots HM, Graybosch RA, Jones H, Parker M, Romanova O, Jones MK, Howe CJ, Trafford K. (2013) Waxy phenotype evolution in the allotetraploid cereal broomcorn millet: mutations at the GBSSI locus in their functional and phylogenetic context. Mol Biol Evol 30:109–122. doi:10.1093/molbev/mss209.

[47] Zi Y, Cheng D, Li H, Guo J, Ju W, Wang C, Humphreys DG, Liu A, Cao X, Liu C, Liu J, Zhao Z, Song J. (2022) Effects of the different waxy proteins on starch biosynthesis, starch physicochemical properties and Chinese noodle quality in wheat. Mol Breed 42:23. doi:10.1007/s11032-022-01292-x.

[48] Benoit M. (1924). Note sur une méthode de résolution des équations normales provenant de l’application de la méthode des moindres carrés a un système d′équations linéaires en nombre inférieur a celui des inconnues. -Application de la méthode a la résolution dún système defini d’équations linéaires (Procédé du commandant Cholesky). Bull. Géodés 2:67–77. doi:10.1007/BF03031308.

[49] Omholt SW, Plahte E, øyehaug L, Xiang K. (2000) Gene regulatory networks generating the phenomena of additivity, dominance and epistasis. Genetics 155:969–980 doi:10.1093/genetics/155.2.969.

[50] de la Fuente A, Snoep JL, Westerhoff HV, Mendes P. (2002) Metabolic control in integrated biochemical systems. Eur J Biochem 269:4399–4408. doi:10.1046/j.1432-1033.2002.03088.x.

