## Supplementary material for "Additive and dominance gene effects described by a logistic function in concentration-limited metabolic pathways at steady state out of equilibrium": Analytical foundations and Figs. S1-S9

### Supplementary Information

#### 1. Relationship between $S_i$ and $dS_i/dE_1$ for a one-step pathway

We investigated the relationship between the variations in substrate ($S_1$) and product ($S_2$) concentrations with respect to the internal enzyme $E_1$—denoted as $dS_1/dE_1$ and $dS_2/dE_1$, respectively—relative to their corresponding steady-state values $S_1$ and $S_2$ for the simplified one-step pathway. The steady-state is given by two conditions: i) mass conservation, i.e. $\sum S_i = S_1 + S_2 = \text{constant}$, and flux balance $J_1 = J_e$. Under these conditions, and according to the Equations (4) and (5) in the main text, we obtain:

$$\frac{E_e k_{cat,e} S_2}{S_2 + K_{M,e}} = \frac{E_1 k_{cat,i} (S_1 - S_2/K_{eq})}{S_1 + K_{M,S_1} (1 + S_2/K_{M,S_2})}. \quad (1)$$

Furthermore, invoking the mass-conservation constraint, Equation (1) can be rewritten as

$$\frac{E_e k_{cat,e} S_2}{S_2 + K_{M,e}} = E_1 k_{cat,i} \frac{((\sum S_i - S_2) - S_2/K_{eq})}{(\sum S_i - S_2) + K_{M,S_1} (1 + S_2/K_{M,S_2})} \quad (2)$$

From (2), $S_2$ is readily obtained in terms of $E_1$ as

$$S_2(E_1) = \frac{\sqrt{K_{M,S_2} \Gamma^2 + \Delta}}{\delta} - \frac{\Gamma}{\delta}, \quad (3)$$

where

$$\Gamma = K_{M,S_2} (k_{cat,e} E_e K_{eq} ((S_1 + S_2) + K_{M,S_1}) + E_1 k_{cat,i} (K_{eq} (K_{M,e} - (S_1 + S_2)) + K_{M,e})),$$

with

$$\Delta = 2(S_1 + S_2) E_1 k_{cat,i} K_{eq} K_{M,e} \delta,$$

and

$$\delta = 2k_{cat,e} E_e K_{eq} (K_{M,S_1} - K_{M,S_2}) + 2E_1 k_{cat,i} (K_{eq} + 1) K_{M,S_2}.$$

To elucidate these ideas, we fixed $\sum S_i = 6 \text{ mM}$ and set $K_{eq} = 10$, $K_{M,S_1} = 0.1 \text{ mM}$, $K_{M,S_2} = 10 \text{ mM}$, $K_{M,e} = 1 \text{ mM}$, $k_{cat,i} = 10000 \text{ min}^{-1}$, $k_{cat,e} = 10000 \text{ min}^{-1}$, and $E_e = 10 \text{ } \mu\text{M}$ which allowed us to establish the relationship $S_2(E_1)$ (3). By a similar procedure we obtained $S_1(E_1)$; both functions are plotted in Fig. S1A,B. It is worth emphasizing that the observed behavior is a direct consequence of assuming that the pathway interfaces with its surroundings via irreversible Michaelis–Menten kinetics (Equation 1).

In Fig. S1B we can appreciate that the function has two asymptotes, $S_i^{\min}$ and $S_i^{\max}$, for each metabolite and that there exists an inflection point between the asymptotes, a behavior that is absent if the input/output enzyme has first-order kinetics as $J_e = kS_2$ with $k = 1000 \text{ min}^{-1}$ (Fig. S1D). Since we considered that coupling to the environment is more realistically represented by an irreversible Michaelis–Menten kinetics (see main text for details), we wanted to obtain more details on the behavior observed in Fig. S1A. To this end, we examined the relationship between the variation in product concentration, denoted as $dS_2/dE_1$, and the concentration $S_2$ itself. For each value of $E_1$, we plot $dS_2/dE_1$ as a function of $S_2$, as shown in Fig. S2.

The profile displayed in Fig. S2 indicates that

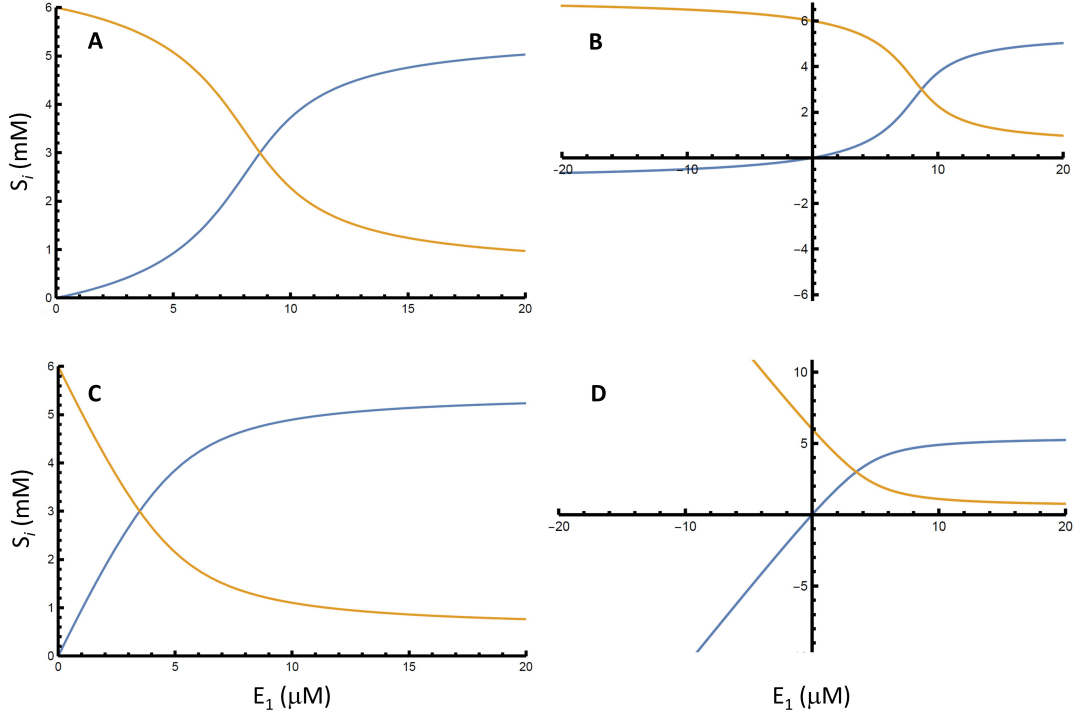

Figure S1: Metabolite concentrations $S_i$ ($S_1$ orange; $S_2$ blue) as a function of enzyme 1 concentration $E_1$ in the one-step pathway calculated from Equation (3) under the assumption that external enzyme has irreversible Michaelis-Menten kinetics (A,B) or first-order kinetics ($J_e = kS_2$) (C,D). In A and C, the plots are shown in the positive quadrant, while in B and D, the same graphs are extended to negative quadrants to better appreciate the asymptotic $S_1^{min}$ and $S_2^{min}$ at minimal $E_1$ when the external enzyme has irreversible Michaelis-Menten kinetics, and the lack of asymptotes at minimal $E_1$ when the external enzyme has first-order kinetics. However, negative quadrants have no physical sense.

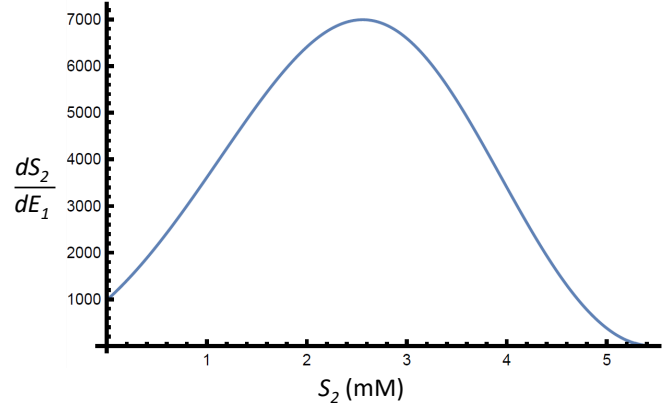

Figure S2: $dS_2/dE_1$ as function of $S_2$, calculated from the plot in Fig. S1A.

$dS_2/dE_1$ varies with $S_2$ according to a sigmoidal pattern:

$$\frac{dS_2(E_1)}{dE_1} = \gamma (C - S_2(E_1)) (S_2(E_1) - D). \quad (4)$$

By symmetry, the preceding analysis can be repeated for $S_1$, leading to an analogous outcome.

The sigmoid behavior presented here—which emerges under the irreversible Michaelis-Menten kinetics for coupling to the environment—was not captured in previous works, since they relied on first-order kinetics for the exchange fluxes (see main text).

### 2. Further details on the $S_i/E_i$ relationship and its coupling to the surroundings

In the main text, the increase of product concentration $S_2$ as a function of $E_1$ for the one-step pathway was calculated using coupled DEs (eq. 8 in the main text) fol-lowing the model described by eq. (6) in the main text, with stationary-state $S_i$. These numerical calculations produced good fit with eq. (12) of the main text, and here we repeated the procedure for substrate $S_1$. Since the reaction is reversible, we expected that the substrate concentration $S_1$ should change similarly as the product $S_2$ shown in Fig. 1, although with opposite slopes. This behavior is confirmed in Fig. S3. Dominance relationships are not shown there because it was assumed that trait $T$ is determined by $S_2$.

In a reinterpretation of additive gene effects we defined the maximal additivity as $A_i = S_i^{max} - S_i^0$. As we pointed out in the main text, this quantity is related with $\gamma_i$, which in turn depends on $E_e$:

$$\gamma_i(E_e) = \frac{\gamma_{min}E_e + \gamma_0E_e^{0.5}}{E_e + E_e^{0.5}}; \quad A = \frac{2(dS_i/dE_1)|_{E_1^0}}{\gamma_i}. \quad (5)$$

Therefore, we wanted to investigate the relationship between these three parameters. As shown in Fig. S4A-C, there is a complex interplay among them. If we center the analysis on the system's coupling to the surroundings as represented by $E_e$, we can see that stronger coupling diminishes both maximal additivity (Fig. S4A) and sensitivity (Fig. S4B). On the other hand, maximal additivity is higher when sensitivity increases (Fig. S4C), in agreement with the trends between these two parameters against $E_e$. The total concentration in the system, expressed as $\sum S_i$, modulates these behaviors, as increments in $\sum S_i$ diminishes maximal additivity (Fig. S4A and C) contrarily to the trend with system's sensitivity (Fig. S4B). In the main text we distinguished environmental influence related with internal conditions (as $\sum S_i$) from that related with external conditions (as $E_e$). Here we can precise more these influences on maximal additivity and observe that internal and external environment move maximal additivity with parallel trends in the sense that when either $\sum S_i$ or $E_e$ increases $A_i$ decreases (Fig. S4A) by difference with sensitivity, which increases with $\sum S_i$ but decreases with $E_e$ (Fig. S4B). Therefore, stronger coupling to the surroundings as well as higher total metabolites concentration diminish the maximal additive gene effects but these effects may possess different rates of response to environmental change.

#### 3. Relationship between the degree of dominance and the enzyme concentrations

We may consider that parent and hybrid offspring phenotypes may be defined by $P_1 = S_2(E_1^{gg})$, $P_2 = S_2(E_1^{GG})$, $H = S_2(E_1^{Gg})$ respectively, being $E_1^{Gg} = (E_1^{GG} + E_1^{gg})/2$. In addition, the mid-point between both parents is $m = S_2(E_1^m)$. Therefore, using the Mean

Value Theorem we can write

$$P_2 - P_1 = \frac{dS_2}{dE_1} \Big|_{E_1^x} (E_1^{GG} - E_1^{gg}), \quad (6)$$

where $E_1^x$ is within the $[E_1^{gg}, E_1^{GG}]$ and $dS_2/dE_1|_{E_1^x}$ is the derivative of the function $S_2$ evaluated at $E_1^x$. Using the definitions (1) from the main text, we have

$$\frac{dS_2}{dE_1} \Big|_{E_1^x} = \frac{P_2 - P_1}{E_1^{GG} - E_1^{gg}} = \frac{2a_2}{E_1^{GG} - E_1^{gg}}. \quad (7)$$

Considering for all practical purposes $E_1^{Gg} \simeq E_1^x$ and $E_1^m \simeq E_1^x$, we can write the linear approximation centered on $E_1^x$

$$\begin{aligned} H = S_2(E_1^{Gg}) &\simeq S_2(E_1^x) + dS_2/dE_1|_{E_1^x} (E_1^{Gg} - E_1^x) = \\ &= S_2(E_1^x) + \frac{2a_2}{E_1^{GG} - E_1^{gg}} (E_1^{Gg} - E_1^x), \end{aligned} \quad (8)$$

$$\begin{aligned} m = S_2(E_1^m) &\simeq S_2(E_1^x) + dS_2/dE_1|_{E_1^x} (E_1^m - E_1^x) = \\ &= S_2(E_1^x) + \frac{2a_2}{E_1^{GG} - E_1^{gg}} (E_1^m - E_1^x). \end{aligned} \quad (9)$$

Using approximations (8) and (9) in definition (1) of the main text:

$$\begin{aligned} d_2 = H - m &\simeq \frac{2a_2(E_1^{Gg} - E_1^m)}{E_1^{GG} - E_1^{gg}} = \\ &= \frac{2a_2E_1^{Gg}(1 - E_1^m/E_1^{Gg})}{E_1^{GG} - E_1^{gg}}. \end{aligned} \quad (10)$$

Considering that $2E_1^{Gg} = (E_1^{GG} + E_1^{gg})$ and therefore $E_1^{GG} - E_1^{gg} = 2E_1^{Gg} - 2E_1^{gg}$, and defining the degree of dominance as $r_2 = d_2/a_2$, the relationship (10) may be written as

$$r_2 = \frac{d_2}{a_2} \simeq \frac{E_1^{Gg} - E_1^m}{E_1^{Gg} - E_1^{gg}}. \quad (11)$$

Thinking of equation (11) as an equality, we can write

$$E_1^m = r_2(E_1^{gg} - E_1^{Gg}) + E_1^{Gg}. \quad (12)$$

#### 4. The $S_i/E_i$ relationships in more complex pathways

We analyzed the logistic curves for pathways larger than one step, and observed that, although the logistic relationship holds for all steps, peaks appeared at steps before the last one, which were explained by adding a Gaussian term to eq. 12. In the main text we showed

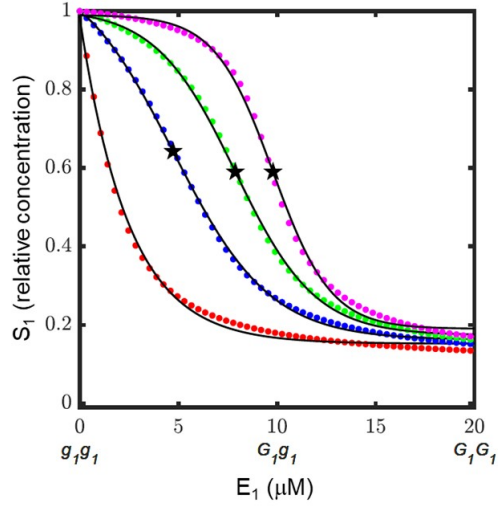

Figure S3: Substrate concentration ($S_1$) in response to increases in active monomeric internal enzyme concentration ($E_1$) in an open system at steady state out of equilibrium ($E_e = 10 \mu M$). Total concentrations $\sum S_i$ were: $0.2 \text{ mM}$ (red), $2.0 \text{ mM}$ (blue), $6.0 \text{ mM}$ (green), and $20.0 \text{ mM}$ (magenta). Black lines are the fitting of data with eq. 12 from the main text ($0.98 \leq R^2 \leq 0.99$), the stars indicating the inflection point (for $\sum S_i = 0.2 \text{ mM}$ inflection point is not shown because it lies at the negative quadrant). Relative $S_1$ concentrations are $S_1/(S_1 + S_2)$ to ease comparison among the different $\sum S_i$.

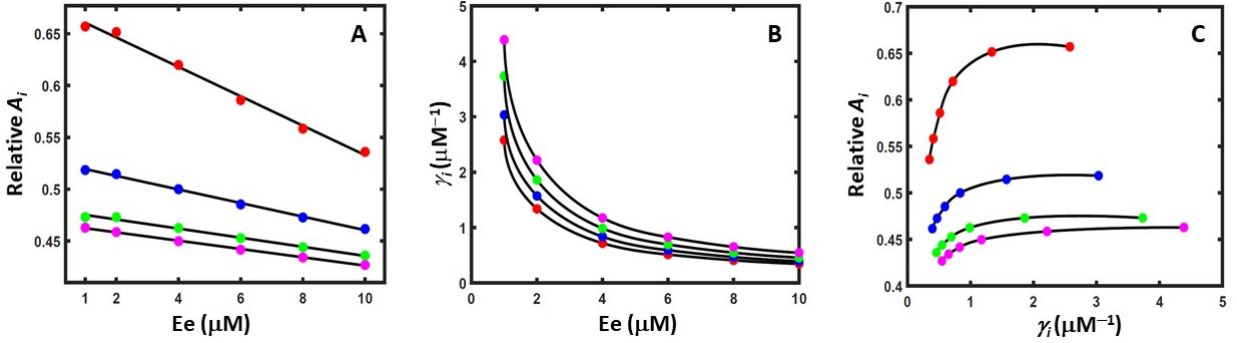

Figure S4: Relationships between $A_i$, $\gamma_i$ and $E_e$, for the same set of $E_e$: 1; 2; 4; 6; 8 and $10 \mu M$. A: $A_i$ against $E_e$; B: $\gamma_i$ against $E_e$; C: $A_i$ against $\gamma_i$. $\sum S_i = 2 \text{ mM}$. (red); $4 \text{ mM}$ (blue); $6 \text{ mM}$ (green); $8 \text{ mM}$ (magenta). Black lines are fits with a linear relationship (A), eq. 5 (B, C); in all cases, ($0.98 \leq R^2 \leq 0.99$). Relative $A_i = (S_i^{max} - S_i^0)/S_i^{max}$.

a chain comprising four reactions where $E_4$ was varied while $E_1$ to $E_3$ were maintained constant, at $5 \mu M$ (Fig. 5 in the main text). We noticed that there was a clear effect of $E_e$: when its concentration was lower than $E_{i \neq 4}$, $S_5^{max}$ accounted for most of the mass of the system, while if $E_e$ was larger than $E_{i \neq 4}$, $S_5^{max}$ was lower than half the mass of the system. Therefore, we investigated these relationships at other $E_{i \neq 4}$, and observed that, indeed, this behavior was the same at $E_{i \neq 4} = 10 \mu M$ or $E_{i \neq 4} = 20 \mu M$ (Fig. S5).

Pathways with different number of steps, from two to six, exhibited the same behavior (Fig. S6). Interestingly, the trends were repeated almost exactly for every reaction regardless of the number of steps, suggesting that local effects are juxtaposed with system-level properties. In addition, the peaks observed in metabolite concentrations $S_{i \neq n}$ at one, two or three reactions before the last one vanished at fourth and fifth reaction before the last one (Fig. S6) indicating that the change of regime pro-voked at the last reaction inflection point did not propagate backwards beyond three reactions.

Moreover, for $S_i$ being the substrate of the third reaction before $E_n$, $E_n^p$ coincides with $E_n^0$, and later, $E_n^p$ tends to diminish linearly as the step into consideration approaches the catalyzed reaction (Fig. S7).

Along this work we assumed that the step affected by the mutation is the last one, but this need not always be the case. Therefore, we also studied the four-step pathway varying either $E_1$, $E_2$ or $E_3$ (Fig. S8). We observed that the pattern observed in Fig. S5 was repeated before the affected step, but later the relationship $S_i(E_i)$ was logistic again, pointing out that the change of regime did not propagate forward. Interestingly, the patterns downstream of the affected enzyme are similar in all cases; however, upstream of it, each pattern seems to be displaced forward in the same direction as the affected step.

The shape of curves obeying eq. 21 of the main text led us to consider that overdominance and dominance re-versals might be explained if we assume that the metabolite responsible for trait $T$ is previous to the substrate or product of the enzyme whose concentration varies among genotypes. In Fig. S9 we show the case in which $T$ is caused by $S_3$ in the four-reaction pathway when $E_4$ is varying. Overdominance may be observed when the hybrid $G_4g_4$, which determines $E_4^{Gg}$ concentration, produces the top of $S_3$ peak, being the contrasting parents at either side (Fig. S9A). Moreover, in Fig. S9B a dominance reversal may be observed for the same metabolite by changing coupling to the environment between strong ($E_e = 10 \mu M$) to middle ($E_e = 6 \mu M$).

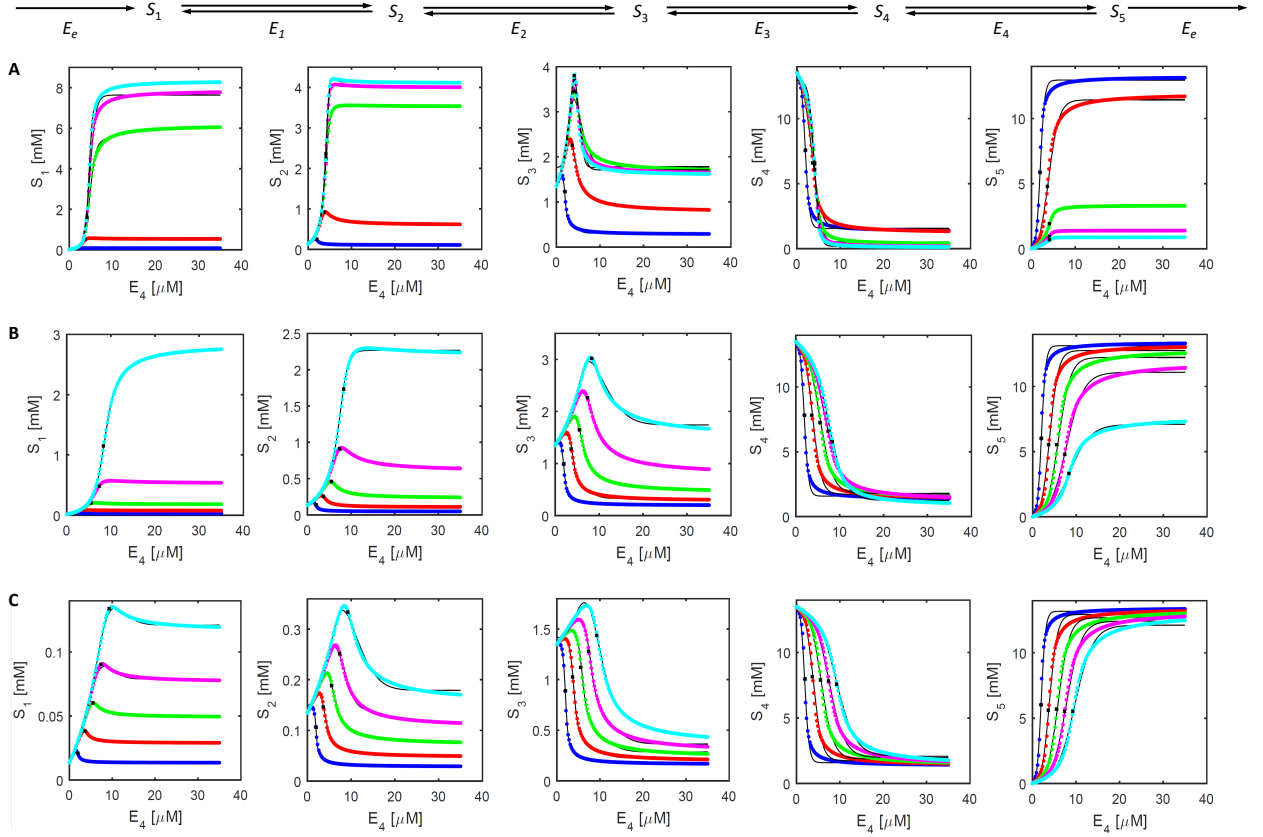

Figure S5: Fit of $S_1$-$S_5$ with eq. 21 of the main text when varying $E_4$. The other enzymes of the pathway, i.e. $E_{i \neq 4}$, were set to $5 \mu M$ (upper panel), $10 \mu M$ (middle panel), and $20 \mu M$ (lower panel). The upper panel reproduces Fig. 5A from the main text, for the purpose of comparison. In all panels, $\sum S_i = 15 \text{ mM}$ since in this condition there is mixed-order reaction regime, and the logistic curve is well defined in the positive quadrant (see Fig. 1 in the main text). In each graph, $E_e = 2 \mu M$ (blue), $4 \mu M$ (red), $6 \mu M$ (green), $8 \mu M$ (magenta), and $10 \mu M$ (cyan), and the curves adjusted according to eq. 21 are shown as continuous black lines ($0.98 \leq R^2 \leq 0.99$). The black asterisk indicates $S_i$ at $E_4^0$ for each metabolite in the pathway.

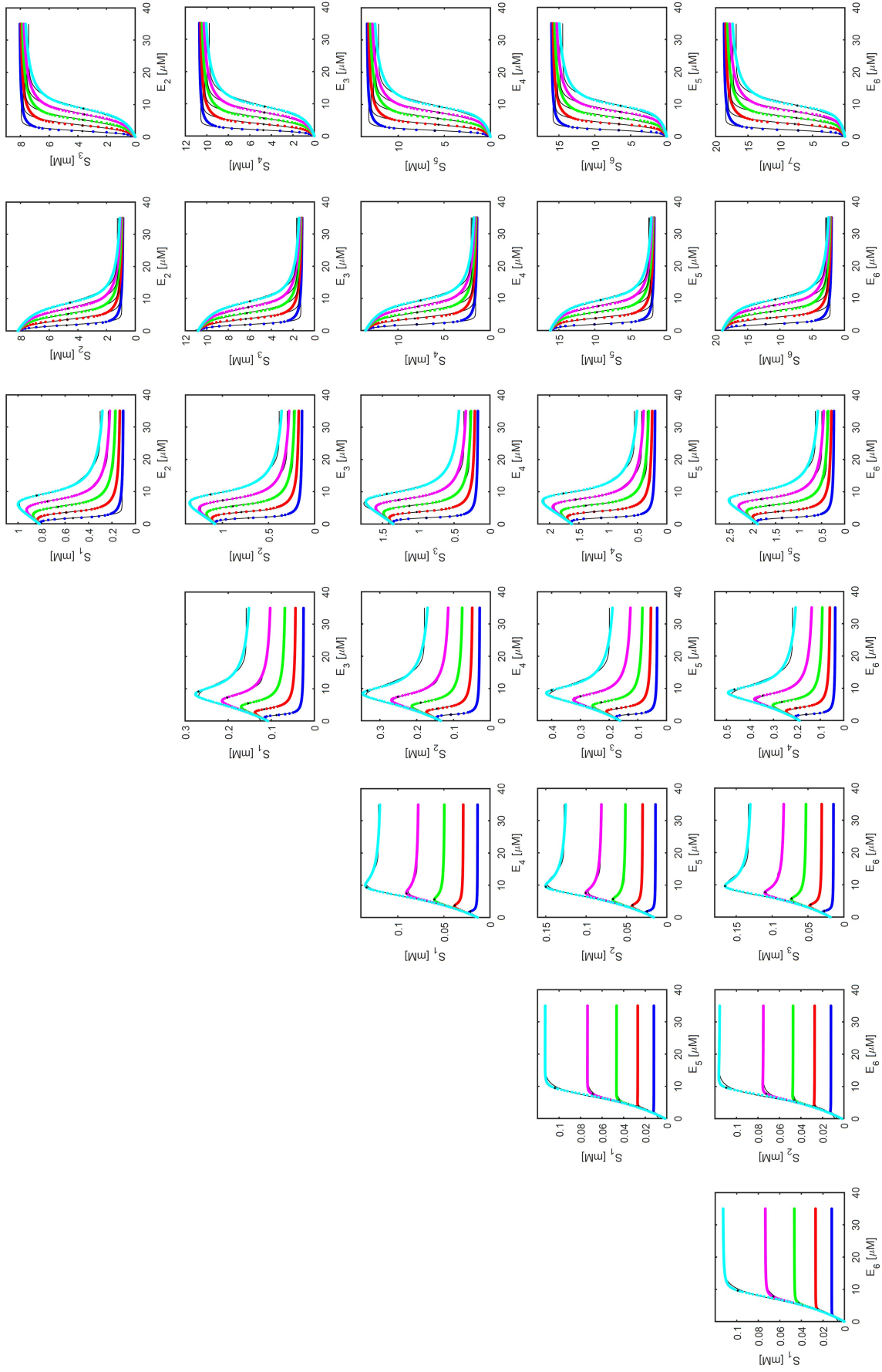

Figure S6: Fit of $S_n$ ($n = 1$ to $7$) with eq. 21 when varying the concentration of the pathway, $E_n$ ($n = 2$ to $6$ from top to bottom). The other enzymes of the pathway were set to $20 \mu M$. The middle panel reproduces the bottom panel from Fig. S5 for better comparison. In all panels, $S_i(0) = 3 mM$ since in this condition there is mixed-order reaction regime, and the logistic curve is well defined in the positive quadrant (see Fig. 1 in the main text). In each graph, $E_e = 2 \mu M$ (blue), $E_e = 4 \mu M$ (red), $E_e = 6 \mu M$ (green), $E_e = 8 \mu M$ (magenta), $E_e = 10 \mu M$ (cyan), and the curves adjusted according to eq. 21 are shown as continuous black lines ($0.98 \leq R^2 \leq 0.99$). The black asterisk indicates $S_i$ at the $E_i^0$ corresponding to the last reaction.

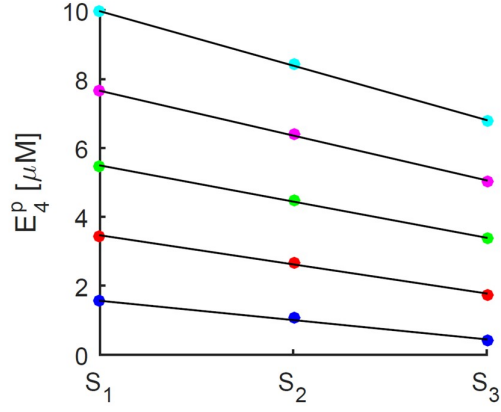

Figure S7: In the four-reaction pathway (middle panel in Fig S6) $E_4^p$ coincides with $E_4^0$ for $S_1$ (values at *ordinates*), and tend to diminish at later steps, for all the connection with the surroundings tested, being them $E_e = 2 \mu M$ (blue), $E_e = 4 \mu M$ (red), $E_e = 6 \mu M$ (green), $E_e = 8 \mu M$ (magenta), $E_e = 10 \mu M$ (cyan). Lines are fits with a linear relationship ($0.98 \leq R^2 \leq 0.99$). As observed in Fig. S6, the peaks are higher as the connection with the surroundings gets stronger. The behavior is repeated in five- and six-reaction pathways (see Fig. S6).

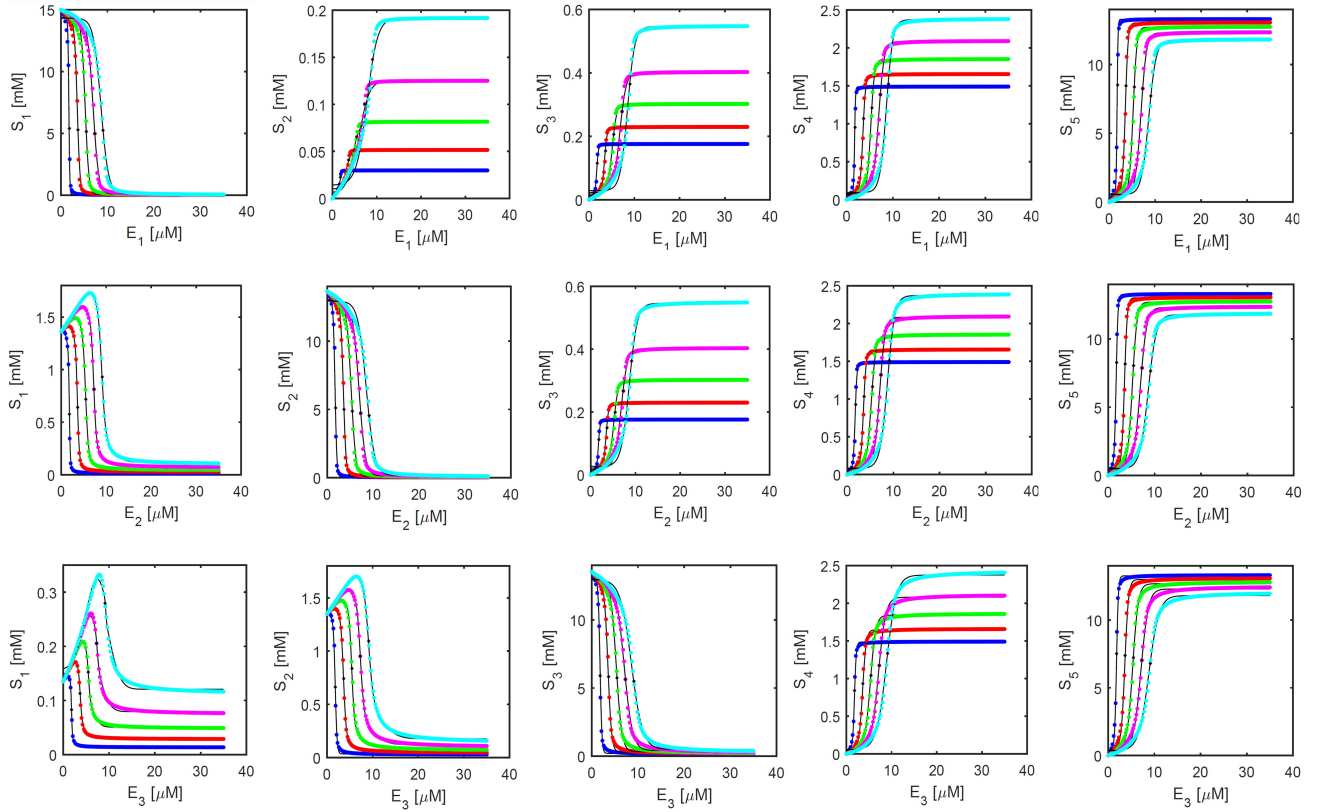

Figure S8: Four-step pathway varying $E_1$ (upper panel), $E_2$ (middle panel) or $E_3$ (lower panel) with $E_e = 2 \mu M$ (blue), $E_e = 4 \mu M$ (red), $E_e = 6 \mu M$ (green), $E_e = 8 \mu M$ (magenta), $E_e = 10 \mu M$ (cyan). Lines are fits with eq. (21) from the main text ($0.98 \leq R^2 \leq 0.99$). The peaks are not observed in the $S_{j>i}$ after the step where $E_i$ was changed, indicating that the change of regime does not propagate forward. In all cases, $\sum S_i = 15 mM$.

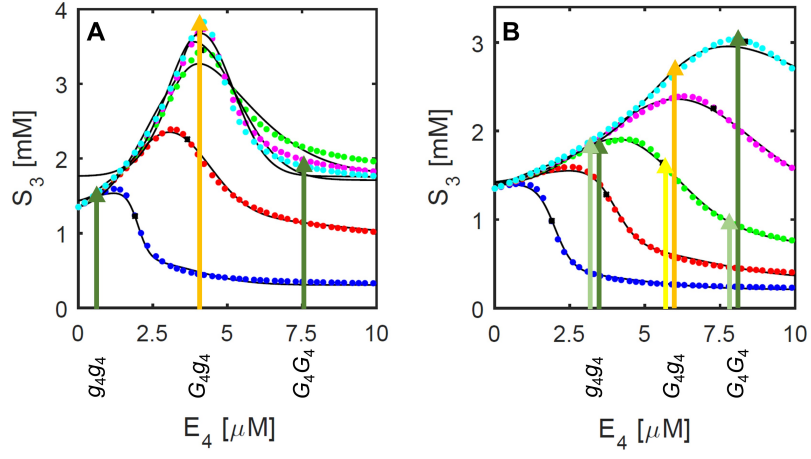

Figure S9: Possible explanations for other dominance relationships. A: overdominance, where the hybrid produces an $E_4^{Gg}$ such that $S_3$ is at the top of its peak (orange arrow), being both homozygous parents at either side, with a small difference in $S_3$ among them (dark green arrows). B: dominance reversal, when coupling to the environment shifts from strong ($E_e = 10 \mu M$; cyan curve, dark color arrows) to middle ($E_e = 6 \mu M$; green curve, light color arrows), which causes inversion of phenotypic values between the homozygous parents. To ease comparison, panels A and B are reproduced from Fig. S5 with $E_{i \neq 4} = 5 \mu M$ (A) and $E_{i \neq 4} = 10 \mu M$ (B), but *abscissa* scales were zoomed in.
